# Spontaneously emerging internal models of visual sequences combine abstract and event-specific information in the prefrontal cortex

**DOI:** 10.1101/2021.10.04.463064

**Authors:** Marie E Bellet, Marion Gay, Joachim Bellet, Bechir Jarraya, Stanislas Dehaene, Timo van Kerkoerle, Theofanis I Panagiotaropoulos

**Author notes:** These authors jointly supervised this work (Stanislas Dehaene, Timo van Kerkoerle, Theofanis I Panagiotaropoulos).

## Abstract

When exposed to sensory sequences, do macaque monkeys spontaneously form abstract internal models that generalize to novel experiences? Here, we show that neuronal populations in macaque ventrolateral prefrontal cortex encode visual sequences by factorizing them into separate codes for the specific pictures presented and for their abstract sequential structure. Ventrolateral prefrontal neurons were recorded while macaque monkeys passively viewed visual sequences and sequence mismatches in the local-global paradigm. Even without any overt task or response requirements, prefrontal populations spontaneously built up representations of sequence structure, serial order, and image identity within distinct but superimposed neuronal subspaces. Representations of sequence structure rapidly updated following single exposure to a mismatch sequence, while orthogonal populations represent mismatches for sequences of different complexity. Finally, those representations generalized across sequences following the same structure but comprising different images. These results suggest that prefrontal populations spontaneously encode rich internal models of visual sequences that reflect both content-specific and abstract information.

## INTRODUCTION

How do we spontaneously encode the specific elements of experiences, and at the same time learn their structure to generalize to new situations? Resolving how the brain encodes sequential patterns of sensory experience on the fly, without any explicit task demands, is key to understand the fundamental computations underlying higher-order cognition. At the neuronal level, internal models implemented by neuronal populations could encode information about a sensory sequence at both an abstract, structural level and at a concrete, sensory level. Importantly, the reinstatement of an internal model, i.e. reactivating the same neuronal ensemble to encode the same structure when it is encountered in new circumstances, may provide a simple mechanism for generalizing knowledge across similar experiences (Bernardi et al., 2020; Chafee and Heilbronner, 2022; Xie et al., 2022). Nevertheless, a complete model should also encode specific aspects of sequential experiences, such as the particular pictures being presented. Therefore, understanding how neuronal populations spontaneously encode sequences of stimuli may provide insights about the abstract scaffolding mechanism that supports generalization, as well as the event-specific neural populations that encode the contents of perception.

Both abstract processing of sequential information and conscious perception of specific stimuli appear to converge in the prefrontal cortex (PFC). PFC is critical for temporal organization of task-related behavior through the encoding, maintenance and flexible use of abstract rules, representations and schemas, that guide behavior and facilitate cognitive control (Miller and Cohen, 2001; Wallis et al., 2001). At the same time, apart from such contextual and abstract information that can be applied to new circumstances, prefrontal populations also encode the specific, consciously perceived contents of sensory experience (Bellet et al., 2022; Kapoor et al., 2022; Panagiotaropoulos et al., 2012). This dual-role of the PFC in representing both abstract and concrete, consciously perceived information suggests that the instantaneous prefrontal population activity should encode both properties, even when a sequence of events is encountered in the absence of any task-related demands. This gives some intuition about the expected structure of the population code during a sequential sensory experience. The neural code should first disentangle, or factorize, the experience into its main structural variables, such as the number of items or the presence of repeated elements (Dehaene et al., 2015; Nieder, 2012; Wang et al., 2015), thus allowing for generalization when similar variables are encountered, a hallmark of abstract processing (Badre et al., 2021; Bernardi et al., 2020). At the same time, apart from abstract information about structure, a complete model should also include concrete information about specific events or contents.

The neuronal properties in the PFC seem to provide the necessary substrate for a population code that can accommodate multiple representations that balance abstract and specific information. In the PFC, a large percentage of neurons exhibit mixed selectivity for the different variables of a task and encode multiple task-related overlapping representations (Fusi et al., 2016; Mante et al., 2013; Rigotti et al., 2013; Saxena and Cunningham, 2019). The mixed selectivity property of prefrontal neurons has the computational capacity to facilitate high-dimensional representations and therefore linear readouts of many different variables from population activity (Badre et al., 2021; Fusi et al., 2016). However, the computational capabilities of PFC would be constrained if the population code allowed only for higher-dimensional representations. In particular, the role of PFC in abstract processing also suggests that prefrontal population activity should also converge into a low-dimensional representation that is optimal for extracting contextual information and generalizing across similar circumstances (Badre et al., 2021; Fusi et al., 2016). With the recent advances in machine-learning decoding techniques, combined with methods that allow recordings of large neuronal populations, it is possible to study how prefrontal population activity encodes these representations.

Here we studied how prefrontal population activity encodes visual sequences and generalizes across them spontaneously, without any overt task or response demands. We used a visual version of the “local-global” paradigm that probes sequence processing at two hierarchical levels, local (sequence element transition probabilities) and global (whole sequence) (Bekinschtein et al., 2009; Chao et al., 2018; Dehaene et al., 2015). We asked two questions. First, do PFC neurons encode all aspects of the sequences in the local-global paradigm, therefore holding a complete internal model of the ongoing sensory stream and its occasional violations? Second, are some of these neural representations abstract enough to be independent of the specific stimulus identities used to convey the sequence pattern, as predicted by earlier work using indirect measures of neuronal activity like functional magnetic resonance imaging (fMRI) (Wang et al., 2015)?

We tested these hypotheses by recording from chronically implanted multielectrode arrays in macaque vlPFC during a local-global paradigm with visual stimuli. Using multivariate decoders, we show that prefrontal neuronal populations spontaneously encoded, within distinct population subspaces, all aspects of the visual sequences, including image identity, serial position of stimuli, and abstract sequence structure, as well as local and global structure violations. These results reveal the fundamental computations in prefrontal ensembles mediating the spontaneous encoding of sequential information.

## RESULTS

### vlPFC spiking activity during the visual local-global paradigm

We recorded spiking activity from neuronal populations with multielectrode Utah arrays, chronically implanted in the vlPFC of two macaque monkeys (Fig. 1A), during exposure to a visual version of the local-global sequence paradigm (Bekinschtein et al., 2009) (Fig. 1B-D). In order to eliminate activity related to decision making and association of sequences with specific motor responses that could confound sequence specific population coding, the task did not require overt behavioral reports, but mere sequence observation. Therefore, this no-report paradigm allowed studying the spontaneous emergence of population codes during passive observation of visual sequences. On each trial, we presented a sequence of four images (300 ms stimulus duration; 300 ms inter-stimulus interval) on a screen while the monkeys maintained their gaze for the whole trial duration within the image region (Fig. 1B). For completion of a trial, the monkeys received a liquid reward 100 ms after offset of a sequence. A sequence consisted either of four repeats of the same stimulus (xxxx sequence, from here on abbreviated as xx) or of 3 repeats and one *local deviant* in the last position (xxxY sequence, abbreviated as xY; Fig. 1C). We presented these sequences in blocks of 200 trials where one sequence type (xx or xY) was the frequent sequence (global standard). The global standard sequence was presented in the first 50 trials of each block. Of the remaining 150 test trials, 80% were global standards and 20% were global deviants, which differed only in the last position compared to the standard. We will use a notation that indicates trials according to their local structure and global context: e.g. a rare xY trial (in an xx block) will be denoted as xY|xx. The first two letters indicate the current trial and the last two letters the global context in which it occurred (Table 1).

**Figure 1.**
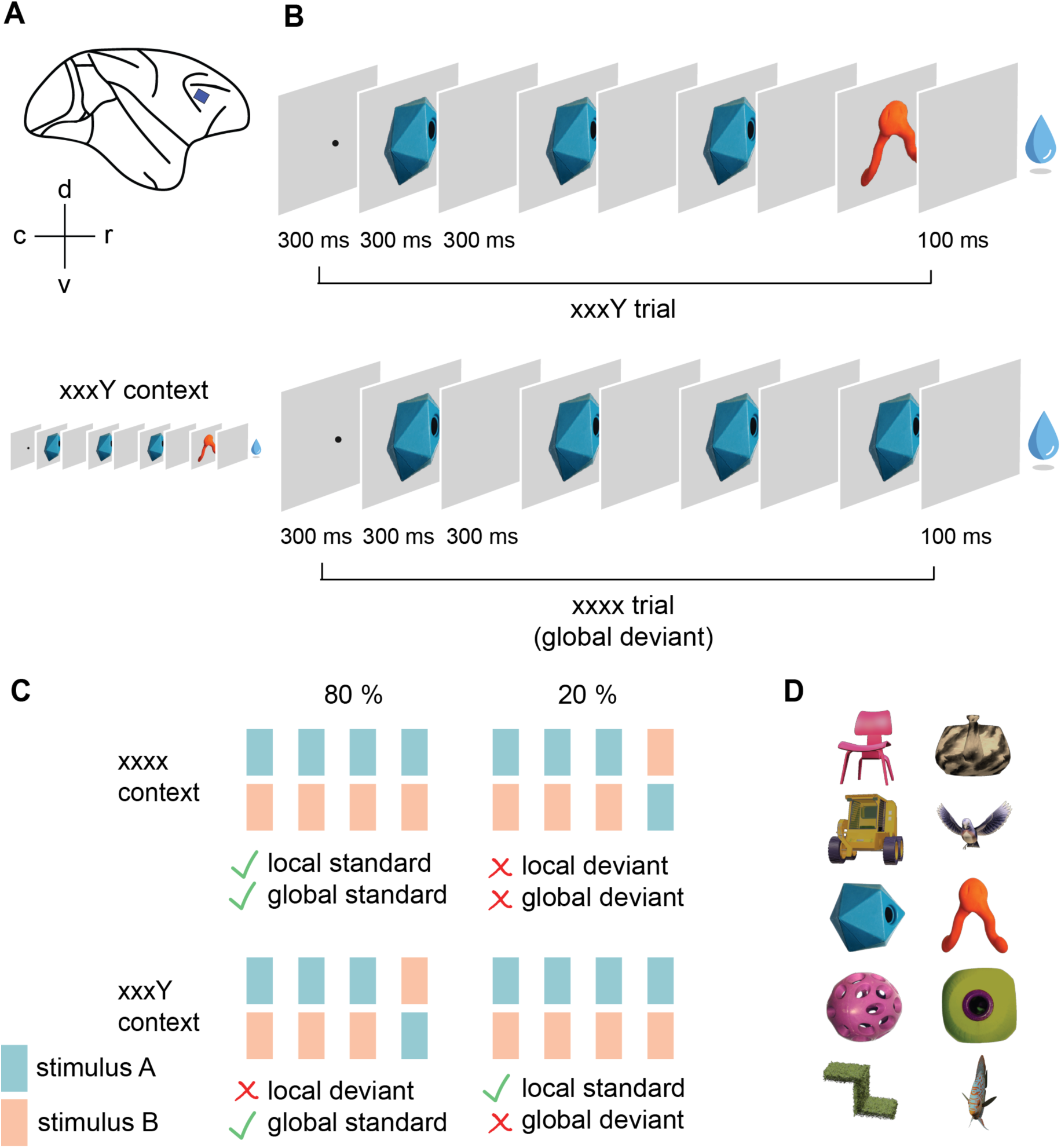
Recording vlPFC spiking activity during the visual local-global paradigm. **A**: Location of the implanted Utah arrays in the macaque vlPFC. **B**: Example trials. On each trial, monkeys fixated for 300 ms prior to sequence onset. A sequence of 4 stimuli was presented with an SOA of 600 ms. 100 ms after offset of the last stimulus, the monkeys received a liquid reward. Examples show a single xxxY and an xxxx trial within the context of frequent xxxY sequences. **C**: Each session consisted of four blocks comprising a frequent sequence (global standard, which could have the structure xxxx or xxxY) and a rare sequence (global deviant). In a given block, the x was a fixed image (A or B, taken from the pairs in panel D) and the Y was the other image (B or A). A block consisted of 200 completed trials. The first 50 trials served as habituation to the global standard sequence (100% of trials), and then we presented a random mixture of 80% global standards and 20% global deviants, which differed only in the identity of the last item. The identity of A and B varied between recording sessions. **D**: The five pairs of visual stimuli (rows) used in the experiments.

**Table 1:** Terminology of sequence types in the local-global paradigm. *The global context corresponds to the structure of the frequent trials in a block. xx is short for a xxxx trial and xY is short for a xxxY trial. Colors indicate the color code used in Figures 2-4.

| Sequence<br>(one trial) | Global<br>context* | Local<br>deviance | Global<br>deviance | Terminology |
| --- | --- | --- | --- | --- |
| xxxx (aaaa, bbbb) | xx block<br>xY block | standard<br>standard | standard<br>deviant | xx\|xx (frequent xx)<br>xx\|xY (rare xx) |
| xxxY (aaaB, bbbA) | xx block<br>xY block | deviant<br>deviant | deviant<br>standard | xY\|xx (rare xY)<br>xY\|xY (frequent xY) |

In each recording session, a specific pair of images (A and B) was chosen, out of five possible pairs (Fig. 1C, D), and four blocks were run with this picture pair, in random order (two xx blocks, aa and bb; and two xY blocks, aB and bA). This allowed us to test whether the population code of sequence structure generalized within sessions, where the order of the images changed, and across sessions, where stimulus identities were different. Furthermore, the design enabled us to distinguish effects of first-order (local) versus higher-order (global) sequence regularity, which require representing the whole sequence pattern (Methods).

The recorded multi-unit activity (MUA), i.e. the sum of recorded spikes from each electrode, was robust over several days in both animals, with overall more active sites in monkey A. First, similar to the classical approach when measuring global brain-imaging signals (Basirat et al., 2014; Bekinschtein et al., 2009; El Karoui et al., 2015; Uhrig et al., 2014), we assessed the univariate responses to local and global violations. Within individual recording sites, we first examined the MUA in response to “pure” local deviance (frequent xY vs. frequent xx trials) and compared them to the effects of “pure global” deviance (rare xx vs. frequent xx). The rank sum statistics provided a measure of the size of the effects of local and global deviance for each channel (Fig. 2A). After controlling for multiple comparisons, about half of the channels revealed a significant response to local deviance, and only ∼ 3 % or 0.8 % to global deviance, in monkey A or H, respectively (Fig. 2B).

**Figure 2.**
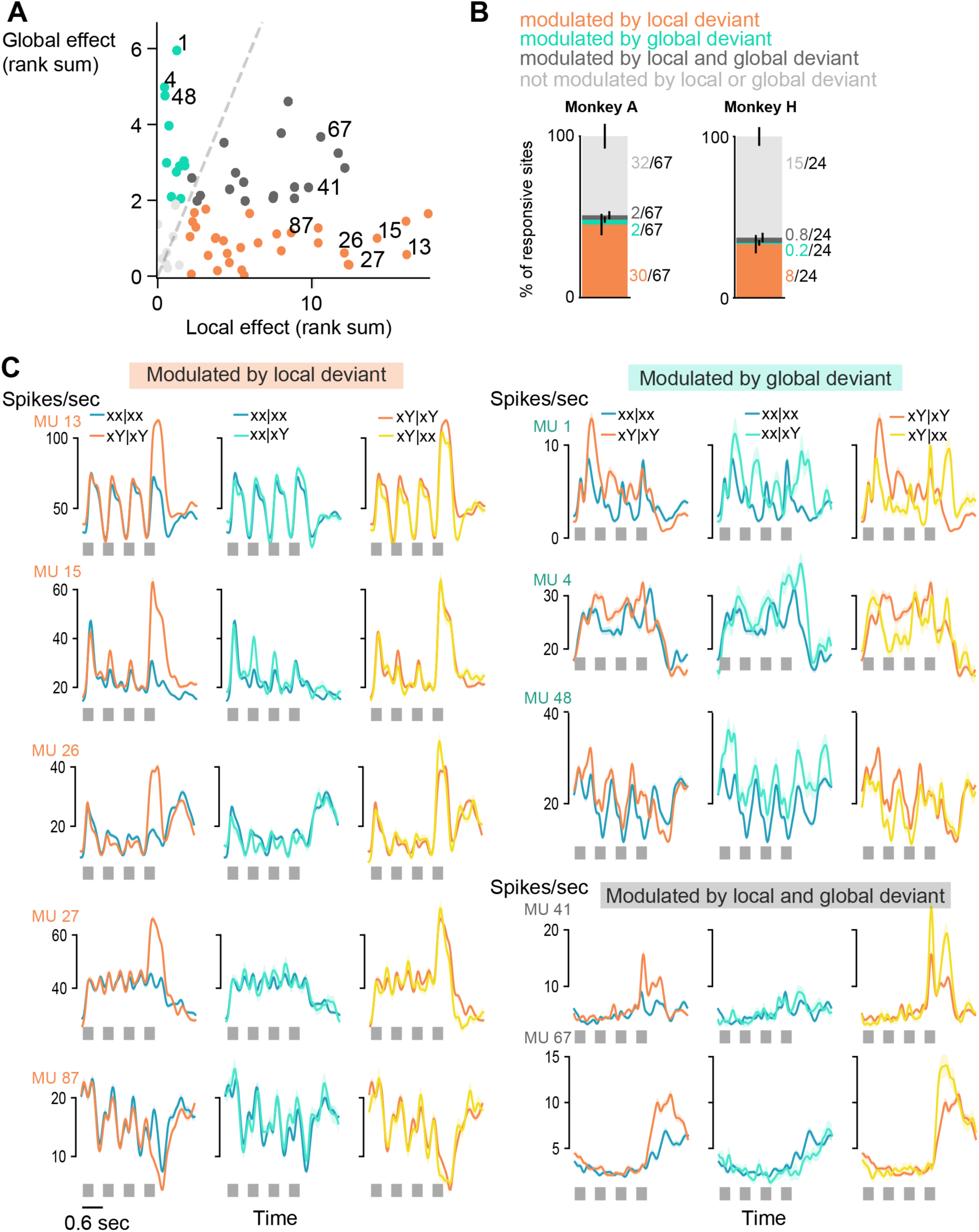
Modulation of spiking activity by local and global sequence structure at individual recording sites. **A**: Effect of local and global deviants (rank sum statistic, p<0.05 prior to correction for multiple comparisons) in task-responsive recording sites (dots) during one session in monkey A. Sites were significantly modulated by local (orange) or global (cyan) deviance only, by both local and global deviance (dark gray) or were not modulated by any deviance (light gray). The dashed gray line indicates equal magnitude of the effect of local and global deviance. **B**: The stacked bar graphs illustrate the proportion of task-responsive sites significantly modulated by local and / or global deviance (67 sites over 6 sessions in monkey A; 24 sites over 10 sessions in monkey H; Mann-Whitney U test with correction for multiple comparisons). Numbers next to the graph correspond to the average number of sites across sessions per condition and error bars (centred on the stacked average proportions across sessions) indicate +-SD of the proportion. **C**: PSTHs of example MUA for sites plotted in **A**. Spiking activity was smoothed using a gaussian kernel with an SD of 50 ms. Colors indicate trial types, averaged over sequences with the A or B image identity. The “pure” local effect is shown by contrasting frequent xx (blue) and frequent xY trials (orange), the “pure” global effect by contrasting between rare vs. frequent xx trials (cyan vs. blue) and the effect of unpredicted over predicted local deviance is shown by contrasting rare vs. frequent xY trials (yellow vs. orange).

The inspection of the peri-stimulus time histogram (PSTH) from individual channels provided some insights into the variety of neural response types (Fig. 2C). Many sites responded strongly to local violations (Fig. 2C left), with a very large firing response to the last image Y in xxxY trials that exceeded the response to any of the previous images. At some, but not all sites, context modulated this response since it was reduced for local deviants that were predicted, i.e. occurring frequently (xY|xY; Fig. 2C, bottom left). Finally, several sites showed distinct firing as a function of global context (xx or xY blocks), already during the first three stimuli of a sequence, and a change in firing when a “pure global deviant” occurred (Fig. 2C, top right).

### VLPFC ensembles form a rich representation of visual sequences

Given the diversity of response types and signs of mixed selectivity, we hypothesized that visual sequences could be better represented in neuronal population vectors rather than within individually specialized neurons, as previously shown for PFC (Badre et al., 2021; Baeg et al., 2003; Ebitz and Hayden, 2021; Fusi et al., 2016; Mante et al., 2013; Parthasarathy et al., 2017; Rigotti et al., 2013). We used regression analyses to test whether vlPFC represented all the variables that defined the visual sequences in the local-global paradigm, namely (i) *stimulus identity* (1 of 2 possible images in each session), (ii) *serial position* of the image within each sequence (from 1 to 4) (iii) *global context* (xx or xY block), (iv) *local deviance*, and (v) *global deviance.* For this analysis, we used only the data from the test trials that followed the first 50 habituation trials in each block, to ensure that the current global context was learned (see also Fig. 4 G,H). We applied multivariate linear regression for all variables but serial position, for which we used multinomial logistic regression (Methods). This approach allowed us to determine which population vectors, if any, carried maximum information about each sequence variable. We then used these vectors to reduce the dimensionality of the MUA and obtain trajectories of the neural population within each neural subspace (Fig. 3). Using the subspace trajectories for classification allowed us to perform a decoding analysis for each of the sequence variables. We quantified such decoding, relative to chance level, using the area under the receiver operating characteristic curve AUROC (see Fig. S1 for Methods).

**Figure 3.**
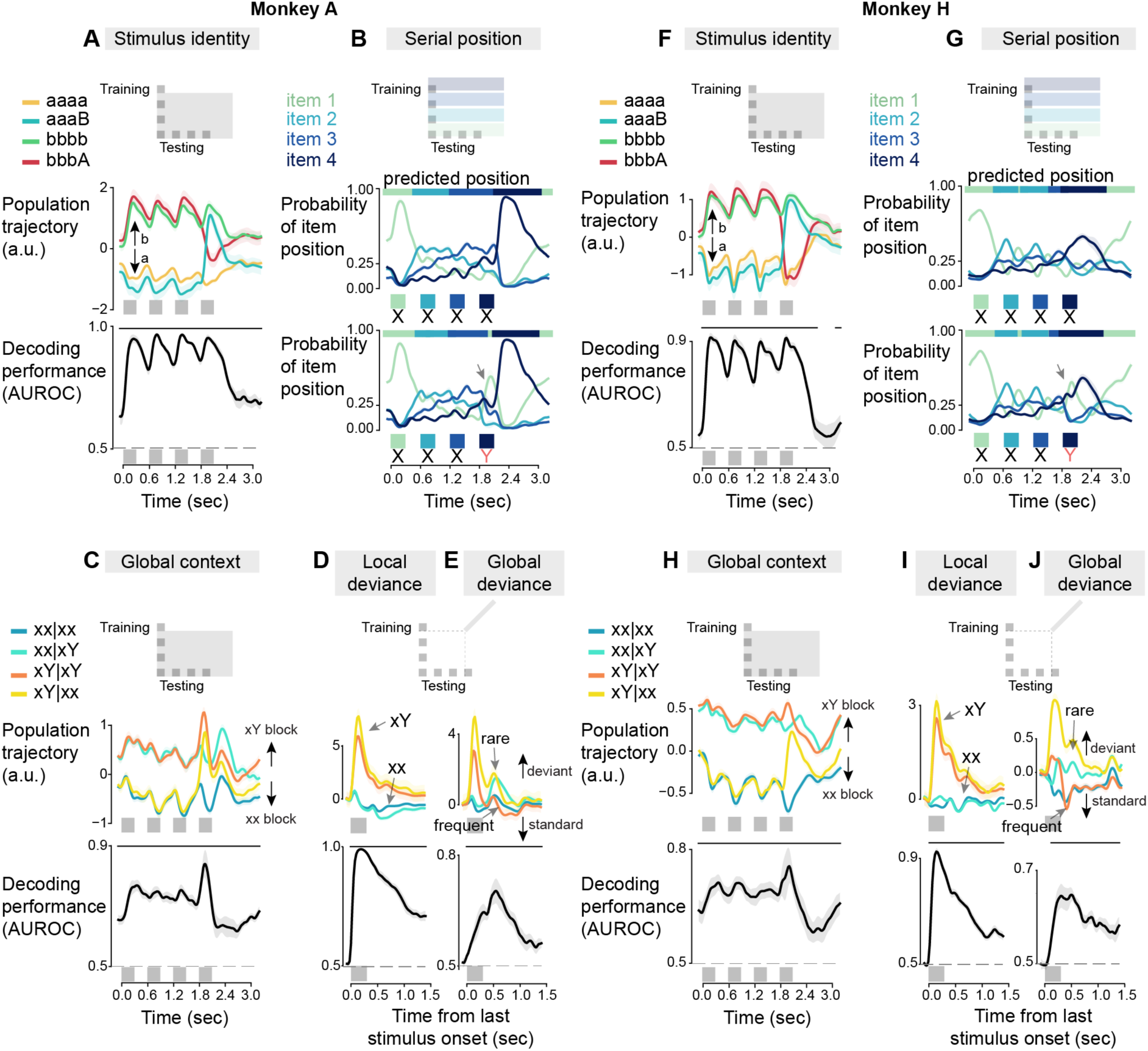
Decoding neural population codes for different aspects of the sequences. Multivariate linear regression was used to estimate the neuronal population vectors encoding stimulus identity, ordinal number, global context, local deviance and global deviance. The upper plots in all panels (and lower plot in B and G) show the population trajectories, i.e. the MUA projected onto the population vectors resulting from the regression (with the time window used for training indicated in the inserts). Black traces in the bottom plots show decoder performance in terms of AUROC, relative to chance level 0.5. Horizontal lines on top of each graph indicate the time points for which the decoding performance was significantly above chance (p<0.05). Data from monkey A (left) and monkey H (right). **A, F**: Decoding of image identity (picture A versus picture B). **B,G**: Decoding of the four ordinal positions in the sequence; the colored curves show the predictive probability of decoders which were trained on xx trials and generalized to xY trials. Note how the population activity in response to the local deviant resembled the response to the first item in a sequence. **C,H**: Decoding of global context, i.e. xY blocks versus xx blocks. Note how the decoding is significant even prior to sequence presentation, indicating an anticipation of the forthcoming sequence). **D,I**: Decoding of local deviance, i.e. xx versus xY sequences. **E,J**: Decoding of global deviance, i.e. frequent versus rare sequences. In panels **D,I,E,J** only, decoding was time-locked to the last sequence item (-100 ms to 1400 ms relative to last stimulus onset). A positive activation indicates a local or global deviance signal, respectively.

**Figure 4.**
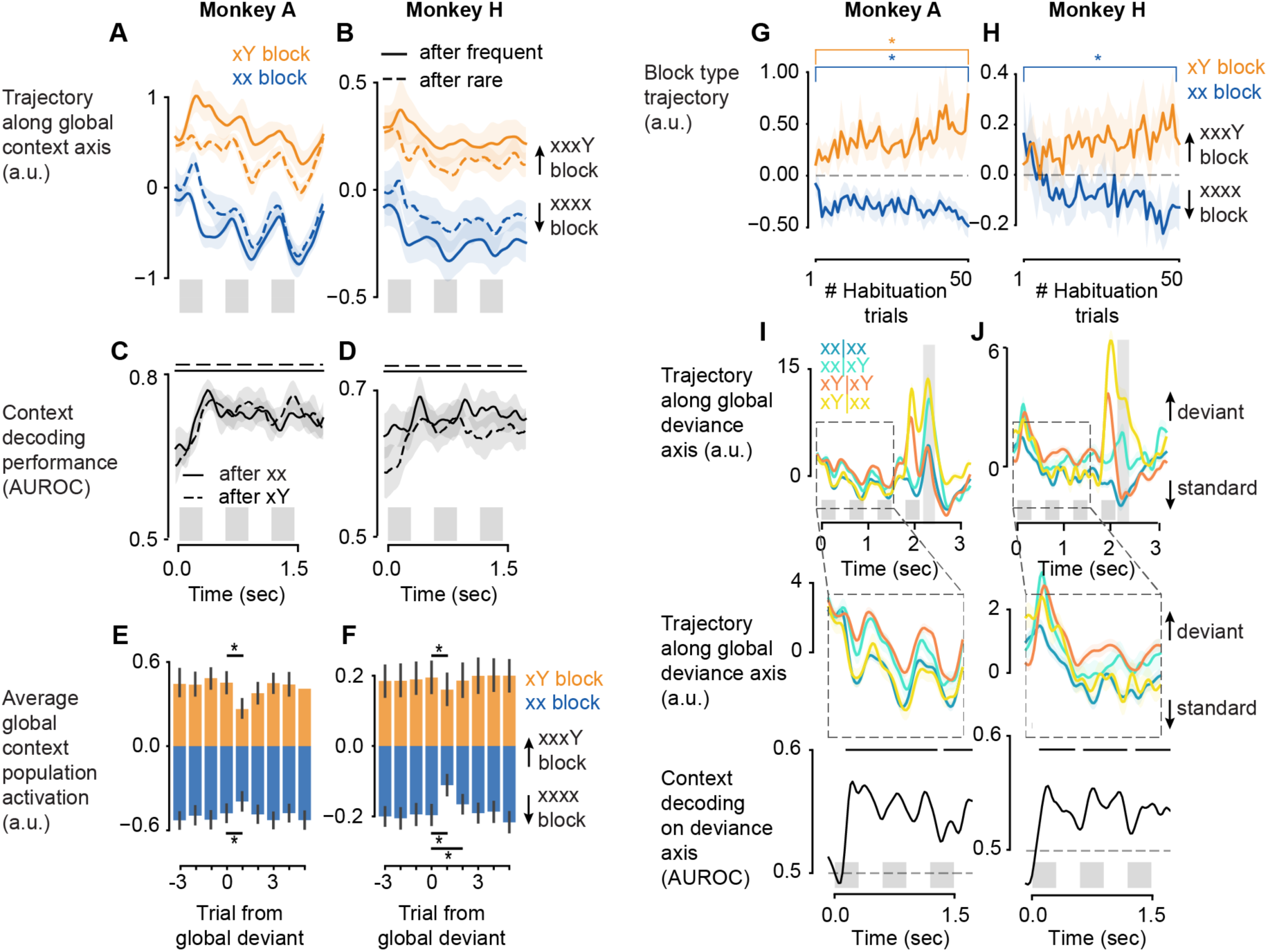
Neural signals reflecting the updating of global sequence knowledge. **A,B**: Population activity projected onto the axis coding for global sequence knowledge, i.e. xY blocks (orange) versus xx blocks (blue). Dashed curves indicate a reduction in the neural separation of xY and xx blocks after a rare global deviant. **C, D**: In both blocks, the global context can still be decoded after a global deviant trial (after xx trials in xY blocks and after xY trials in xx blocks), i.e. when the previous trial is suggestive of the opposite block. **E, F**: Activation of the global context axis averaged over the first three stimuli in each trial, aligned to rare trials in xY blocks (orange) or xx blocks (blue). Height of bars indicate average across trials from pooled sessions and error bars are the 95% CI. The asterisks denote a significant change in global context signal between the trial before the global deviance occurred (0) and the following trial (p<0.05, paired t-test). **G, H**: Buildup of sequence knowledge during habituation. The population activity during the 50 habituation trials of each block was projected onto the population axis that encoded global context and averaged over the first three stimuli in each trial. The asterisk denotes a significant difference (p<0.05) between the first and last habituation trial in a paired t-test across 14 blocks in monkey A and 20 blocks in monkey H (i.e. two blocks of each type per session). **I, J**: Trajectories of the neural population that leads to maximal overall global deviance decoding performance (Fig. 3E,J), indicated by the gray shaded area. The middle panel zooms into the time period prior to the onset of the last stimulus and the bottom panel is the quantification of the context-decoding performance based on the activation of the global deviance population. The horizontal bars on top of the AUROC plots indicate periods of significant context decoding. The horizontal dashed line shows the chance level. Only trials following a global standard sequence were used. Curves indicate average across all trials from all sessions and shaded areas are +-sem.

The results showed that all sequence variables could be decoded at above-chance levels (Fig. 3). First, the decoding performance for stimulus identity (image A vs. B within each recording session) was close to 1 for every item in a sequence, including the last sequence item when it changed on xY trials (Fig. 3A,F). Thus, vlPFC populations robustly encoded the image identity throughout the entire trial.

Second, using a separate decoder for serial position within a sequence, we could predict serial position from neuronal population activity, particularly for the first and last items, reflecting known primacy and recency effects (Chen et al., 1997; Orlov et al., 2000; Terrace et al., 2003), but also at above-chance levels for positions 2 and 3 (Fig. 3B,F, predictions are indicated by horizontal bars). Note that, because the sequences used a fixed timing, this decoding could reflect numerical codes, temporal codes or both (Kapoor et al., 2018; Nieder, 2012). However, elapsed time alone could not explain all of the findings, such as the fact that the code for “1^st^ item” was partially reactivated for the last image of xY trials (the first image with this identity); or that the code for “4^th^ item” was reactivated at ordinal positions 1, 2 or 3 on trials when the monkey broke fixation and the visual sequence was aborted, suggesting that it actually responded to “last item” (Fig. S2). These findings indicate that those population codes were partially locked to the phase of the task, and not solely to timing.

Third, to test whether vlPFC neurons contained a model of the upcoming sequence structure, we decoded the global context (xx or xY block) from the neuronal population activity prior to the last stimulus of a sequence. We indeed identified a population subspace whose activity allowed us to infer the global context even before the sequence presentation started (Fig. 3C,H). Thus, vlPFC populations represented the sequence that recurred in a given block prior to sequence presentation, and not just prior to or after the local deviant. In the following section, we will look in more detail at the properties of this neural subspace and how it builds up during the habituation period.

Fourth, we assessed the presence of responses to violations of either local or global sequence regularity. The population that was sensitive to local deviance showed a response to both predicted (xY|xY) and unpredicted (xY|xx) local deviants (Fig. 3D, I orange and yellow). Decoding local deviance was almost perfect on a single-trial basis, with an early peak (∼200 ms), indicating a very robust and fast response to local novelty. Nevertheless, the activation was stronger upon unpredicted than predicted local deviants, in agreement with the predictive coding framework (Fig. S4). We could also decode global deviance (rare versus frequent sequences in a given block) showing a later peak ∼500 ms after last item onset. In contrast to the unimodal phasic response observed for local deviants, the trajectories of the population encoding global deviance showed a bi-phasic response. First, until around 300 ms (200 ms in monkey H) following the last stimulus onset, only trials with a local deviant showed a positive activation (Fig. 3E,J orange and yellow), again with a higher amplitude for unpredicted local deviants. However, in a later phase, after 300ms, both types of rare trials (xY|xx and xx|xY) evoked a response into the same direction (yellow and cyan), and opposite to the frequent trials (orange and blue). We separately measured the global deviance decoding performance by computing the AUROC for rare vs. frequent xY (Fig. S3 A,B dashed) and rare vs. frequent xx trials (Fig. S3 A,B solid). This analysis confirmed that an early response to global deviants was only present for rare xY trials, while a later mismatch response was present for both sequence types, thus reflecting abstract global deviance detection, invariant for sequence pattern.

We further probed the temporal overlap of these abstract representations, by evaluating a “cross-condition” decoder trained on xx trials and tested on xY trials (Fig. S3 C,D). Generalization across the two sequence structures only occurred late after the last item onset (∼300-500 ms), indicating that, by that time, the neural code for global surprise was shared by the two sequences with a different local structure. This finding also suggests that the population subspace coding for global deviance might be different from the respective subspace coding earlier for local deviance. We further assessed the independence of these two subspaces by decoding global deviance from the subspace that represented local deviance and found that the late global mismatch response was indeed not encoded in the subspace that showed a late local mismatch response (Fig. S4).

Importantly, the generation of eye movements, which were found to be a behavioral read-out of local novelty detection, could not explain neural deviance responses, as the effects remained after the removal of trials in which the monkeys made an eye movement (Fig. S5). The different time scales of the local and global effect are consistent with previous results, showing that the processing of global sequence violations requires longer times that reflect conscious integration compared to the detection of local deviants that can happen non consciously ^32,41,42^.

Altogether, these findings indicate that the vlPFC population that we sampled comprised multiple, overlapping and distributed representations of all the features of the visual sequences used in the local global paradigm.

### The representation of global context is learned spontaneously during habituation and is updated upon errors

Predictive coding models suggest that, following a global context violation, the internal model is destabilized or updated, at least transiently. We examined whether such an update could be detected in the population activity within the subspace that represented global context (xx or xY sequence). Thus, Figure 4 A, B shows population activity projected on the global context axis, depending on both the context (xx or xY) and the preceding trial (global standard or global deviant), prior to and during the presentation of the first 3 sequence items. In both monkeys, there was sustained activity persisting throughout all trials and distinguishing xx from xY context, regardless of the previous trial (Fig. 4C, D). A trial in an xY block, for example, led to an activation into the “xY-direction” of this subspace during the first three stimuli in a sequence, whether following a global standard xY|xY (Fig. 4A,B orange solid) or a global deviant xx|xY trial (Fig. 4A,B orange dashed). This observation is important as it indicates that the activity was not simply due to a lingering memory of the previous trial, but was sustained in the long term, as needed to encode the global context of an entire session. Nevertheless, the encoding strength of global context was reduced after the occurrence of a global deviant (Fig. 4 A,B dashed vs. solid). This effect was transient, and activity was quickly restored within the following 1-2 standard trials (Fig. 4E,F). We also examined how fast this activity was spontaneously built up during the first 50 habituation trials of a block (Fig. 4G,H). The divergence between xx and xY blocks was present early on, and definitely within the first 10 presentations of a given sequence, but it continued to increase continuously through the habituation period of 50 trials.

Together, these findings show that the population activity reflected the global sequence context that was inferred at a long time scale, while at the same time being updated, on a shorter time scale, whenever a deviant sequence occurred. The former finding indicates the spontaneous emergence of a neural representation of global sequence regularity in vlPFC, while the latter fits with a transient destabilization or update of this model after a mismatch to the learned global structure.

### A shared population code for global deviance and context

Current models of predictive coding hypothesize that, at each level of the cortical hierarchy, separate neural populations code for predictions and prediction errors (for a review see Walsh et al., 2020). According to this hypothesis, expectation and mismatch or error signals should be detected in distinct, segregated ensembles. Alternatively, the representation of context might happen within the same neural population that also emits the mismatch responses. Such an integration might be especially relevant for higher-order areas such as PFC, where mismatch signals interact with model representations to promote model update. To address this issue, we studied the overlap of deviant and context representations in PFC. We projected the unfolding population MUA onto the vector corresponding to maximal global deviance decoding performance (+-100 ms around the maximum time bin) (Fig. 4I, J). We found that the resulting population trajectories also segregated as a function of the global context, i.e. xx versus xY blocks (Fig. 4I,J). During the first three items, there was a slightly larger activation into the direction of global deviance on xY blocks than on xx blocks. This was reflected by a significant AUROC for context decoding from these trajectories (Fig. 4I,J bottom). It is important to note that, conversely, this effect of context cannot entirely explain the deviance response to rare xx trials, because the latter elicited an additional increase in activity after the last stimulus (Fig. 4 I,J top cyan). We therefore conclude that at least part of the population code for global deviance detection was shared with the representation of global context.

### Population code for sequence structure generalizes across stimulus identities

The above results establish that vlPFC population activity spontaneously forms partially overlapping representations of for image identity, serial order, global context, local deviance, and global deviance. Furthermore, the above analyses were applied across the two pictures that were presented in a given session (A and B), hinting at the existence of neural codes that generalize across stimulus identities. We exploited the chronic nature of recordings and tested the generalization of the population codes across sessions with different picture pairs, always presented on different days. We found that the same neuronal population vectors allowed us to decode global sequence context, local deviants, and global deviants, respectively, even for stimulus pairs that differed completely from those presented in the training session (Fig. 5). This finding indicates that vlPFC sequence representations were stable for multiple days and, most importantly, were reinstated across distinct picture identities, thus possibly reflecting an abstract neural code. We should note that, although the decoding performance was significantly above chance levels, it was not perfect. This finding may indicate that part of the representation for sequence structure is sequence specific, or it may reflect instabilities in recording from the same neuronal populations across days.

**Figure 5.**
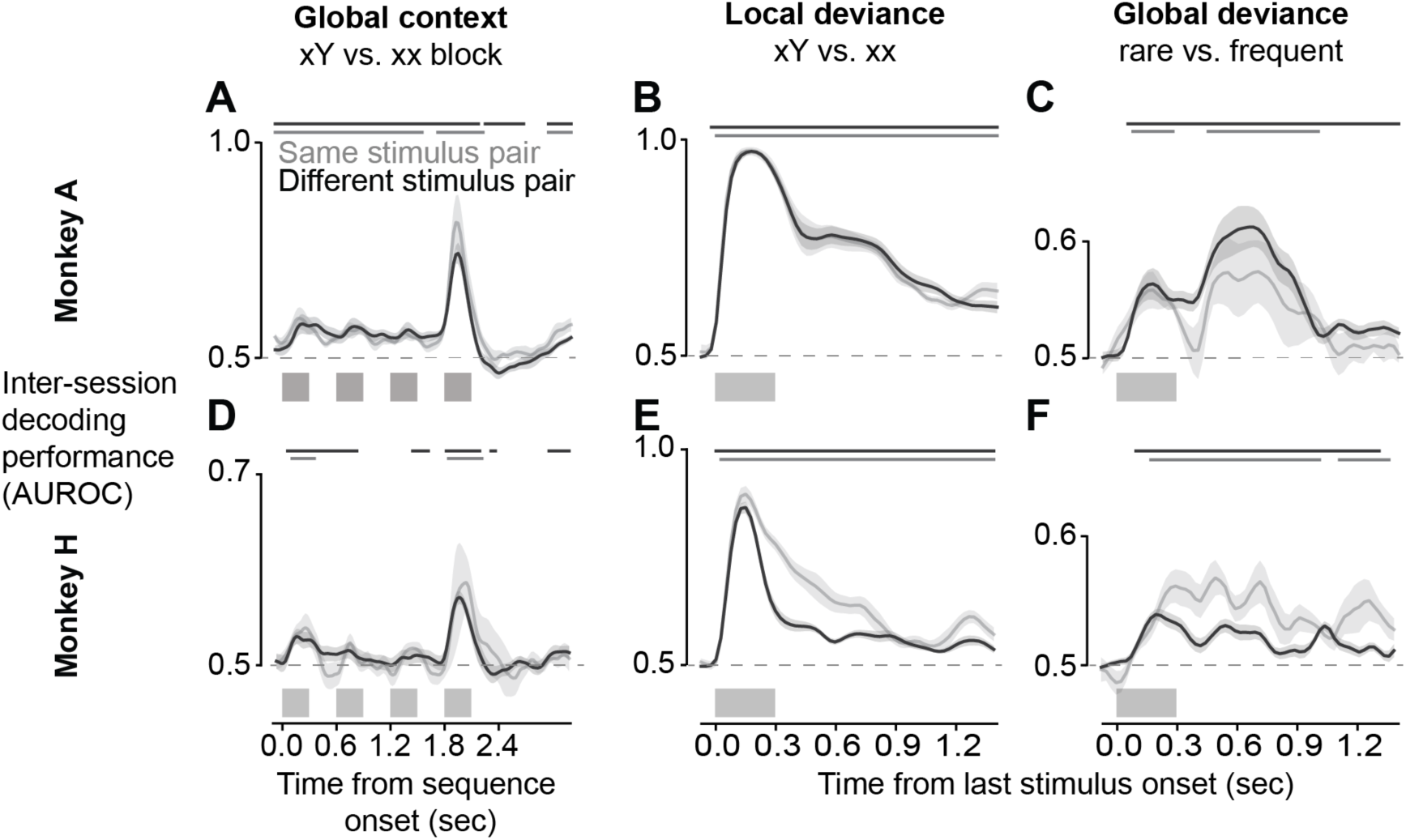
Population responses to global context, local deviance, and global deviance generalize to new sessions and stimuli. Plots show the generalization of decoding to new sessions with either the same visual stimuli (gray curve) or difference stimuli (black curve). Generalization performance (AUROC) is shown separately for decoders trained on global context (**A,D**), local deviance (**B,E**) and global deviance (**C,F**). Horizontal lines on top of each graph show time points where the decoding performance was significantly above chance (p<0.05).

Rather than a difference between predicted and seen pictures, the representation of local deviance could reflect the indirect effect of stimulus-specific adaptation (SSA) occurring at an earlier stage such as inferotemporal cortex (IT). Neural responses would be smaller on BBBB trials than on AAAB trials because the response to picture B would have been adapted (Garrido et al., 2009; May and Tiitinen, 2010). Note that SSA in IT is picture-specific (Meyer and Olson, 2011), but if the signal from multiple picture-specific neurons in IT was integrated in vlPFC, it would explain the observed generalization across different pictures. To test whether the vlPFC population response to local deviants reflected genuine deviance detection or merely SSA, we performed an additional experiment in monkey A, presenting new random sequences with different numbers of image repetitions and changes (Figure 6A, letters indicate any of >900 images). Contrasting the activation evoked by the last stimulus in XXXY sequences with the last stimulus in WXYZ sequences allowed us to disentangle deviance and SSA, as done with the many-standards control in mismatch negativity (MMN) studies (Ruhnau et al., 2012). In the many-standards control paradigm deviance detection predicts a novelty response to XXXY (where a prediction develops about X), but none to the unpredictable sequence WXYZ, as the image identity of each item changed on every trial; SSA, however predicts no difference, as the last picture is equally novel and non-adapted in both cases. The results supported deviance detection (see direction of activation in Fig. 6A and decoding performance in Fig. 6B): local deviants always led to a larger response of this neural subpopulation (Fig. 6C; Fig. S6) indicating that vlPFC indeed encodes deviance from a local context, regardless of picture identity. Indeed, we found that MUA responses underlying this population subspace were far more diverse than what would be expected based on the adaptation hypothesis (Fig. 6D), namely a simple decrease in response amplitude upon repetition. This provides additional evidence that PFC populations spontaneously encode sequence deviance in an abstract way: as previously inferred indirectly through brain-imaging signals (Wang et al., 2015), a population of PFC neurons signals that “the last sequence item differed from the preceding ones”.

**Figure 6.**
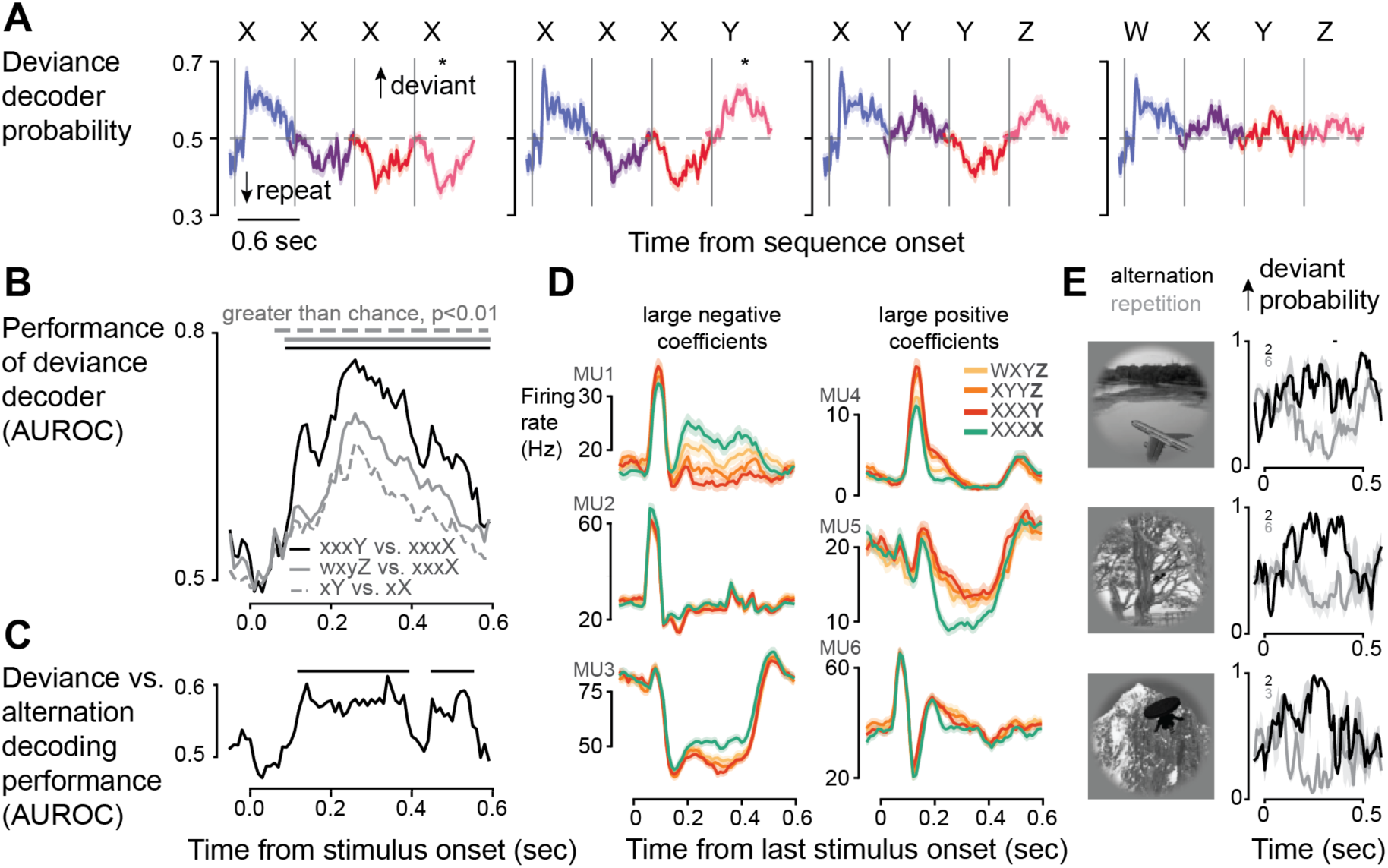
Abstract change and deviance detection by neural populations in monkey A. In a control experiment in monkey A, four possible sequence chunks (see titles in **A**) were presented in a uniform random manner. Letters W-Z indicate any of 948 grayscale images from the Brainscore database (Majaj et al., 2015), changing randomly in each trial. A decoder for local deviance was trained on XXXY vs. XXXX trials, indicated by the star in **A**, using leave-one-stimulus-out cross-validation. **A**: Predictive probability of the decoder for all sequence types and stimulus positions in a sequence (indicated by colors). **B**: Decoding performance in terms of AUROC for local deviants (XXXY vs. XXXX, black), novel stimulus, as well versus pure repeats (WXYZ vs. XXXX, gray), or any transition versus single repeats (XY vs. XX, dashed gray). All conditions could be decoded above chance level with p<0.01 (random permutation test). Horizontal bars indicate significant time bins. **C**: Rare stimuli that violate a local pattern of repetitions (XXXY) yielded a significantly higher response of this population than rare stimuli without preceding regularity (WXYZ), again indicated by horizontal bars on top. **D**: Examples of multi-units contributing to the population axis coding for local deviance. A large negative (left) or positive (right) coefficient means a lower, or higher firing rate for deviant stimuli, respectively. **E**: Population deviance response to repeats (light gray) or alternations (black) of three example images. Lines show mean across N number of trials (indicated by small numbers in each plot) and shaded areas show +-sem.

## DISCUSSION

Our results suggest that prefrontal population activity encodes concrete, event-specific features of sequential information like image identity, but also all other abstract aspects of visual sequences, such as the serial position of each picture and/or task phase, the global sequence pattern, as well as local and global sequence structure violations. The population code for global sequence pattern generalized across similar pattern sequences, suggesting that it encoded abstract structural knowledge. Importantly, these representations emerged spontaneously during passive viewing, in the absence of any active task that could have resulted in the association of specific sequences with behavioral responses or other contingencies, therefore conflating decoding results.

### Abstract sequence processing in the PFC

Single neurons in the PFC encode abstract information about stimulus category (Freedman et al., 2001), number (Nieder et al., 2002) and rules (Wallis et al., 2001). More recently, the development of multichannel recording techniques and machine learning analytical tools, has allowed for the sampling and analysis of the activity of large neuronal populations. This resulted in the reemergence of the Hebbian idea (Hebb, 2005) that joint neuronal population activity is the basic computation unit in the brain (Ebitz and Hayden, 2021; Saxena and Cunningham, 2019; Yuste, 2015). Abstract coding in the PFC has been recently probed at this level of ensemble activity. Population activity in two frontal areas, the orbitofrontal and anterior cingulate cortex, was shown to encode abstract information in a task where stimulus identification and knowledge of context were both necessary to predict reward (Saez et al., 2015). Population activity in the PFC (but also in the hippocampus) was shown to have a low-dimensional geometrical structure that allowed the linear classification of abstract features defined by the association of a stimulus with its operant and reinforcement contingencies (Bernardi et al., 2020). At the same time the code was high-dimensional enough to allow the classification of diverse combinations of specific inputs. The same geometrical properties exist in the human medial frontal cortex for cognitive control, where the population code was both task-general for performance monitoring signals and task-specific in two paradigms, the Stroop task and a multisource interference task (Fu et al., 2022).

Our results demonstrate the ability of the prefrontal population code to represent simultaneously such abstract and task-specific variables spontaneously, through habituation, in a no report sequence paradigm. We suggest that this mechanism could be the basic computational unit mediating the perception of sequences. Learning of abstract features can occur spontaneously through observation, without specific instructions or associations of stimuli with rewards or other contingencies. The population code in the PFC could therefore spontaneously extract abstract properties of sequential information and at the same time encode specific variables like the content of these experiences.

### Implications for predictive processing

Within the theoretical framework of predictive processing, we can conceptualize these prefrontal representations of sequences as internal mental models that may also operate to generate predictions onto the external world (Euler, 2018; Meirhaeghe et al., 2021; Pinotsis et al., 2019; Summerfield et al., 2006, 2008a). The implementation of such mental models and predictive coding computations has not been yet demonstrated at the PFC cell assembly level, but only inferred indirectly from macroscopic brain signals related to sequence violations (Chao et al., 2018; Gil-da-Costa et al., 2013; Uhrig et al., 2014; Wang et al., 2015; Wilson et al., 2017). Here we show that even under passive sensory stimulation conditions, assemblies of PFC neurons form abstract mental models of statistical regularities that are frequently encountered in the sensory input. The function of these models could be to project predictions that bias perception, guide decisions and facilitate sensory processing (Dehaene et al., 2015; Friston, 2005; Summerfield and de Lange, 2014).

In such predictive-coding models, the incongruence of sensory information with an already established mental model results in a surprise signal that serves to update the model (Friston, 2005). Based on this main hypothesis of predictive coding, deviations from learned stimulus sequences of variable complexity have been used to probe the respective complexity of predictive representations in the brain, showing that these models unfold at multiple hierarchical scales, reflecting different levels of abstraction and engaging different cortical areas (Dehaene et al., 2015).

The most commonly studied brain signal to sequence deviants is the mismatch negativity (MMN; Näätänen et al., 1978). In the classical auditory oddball paradigm, a local regularity is established by the repetition of one tone, which leads to a reduction in brain responses; at this point, the presentation of a rare (deviant) tone yields the MMN signal, a stronger negative deflection of the scalp EEG compared to repeated stimuli (Näätänen et al., 2007). Although feed-forward adaptation may contribute to the MMN (Garrido et al., 2009; May and Tiitinen, 2010), several results indicate that it, and similar visual violation responses (Pazo-Alvarez et al., 2003) primarily reflects predictive processing (Garrido et al., 2009; Summerfield et al., 2008a; Wacongne et al., 2012; Winkler, 2007). The source of auditory MMN has been localized in the secondary auditory cortex, superior temporal gyrus and PFC (Deouell, 2007; Dürschmid et al., 2016; Shalgi and Deouell, 2007).

The primate brain does not stop at encoding repetitions like in the MMN. Multiple paradigms indicate that both humans and non-human primates can infer sequence regularities at a more temporally extended and abstract level, for instance the extraction of the same structure or pattern from sequences comprised from different stimuli (e.g. AAB and CCD) (Wang et al., 2015). Processing such higher-order sequential structures necessarily abstracts away from the specific stimulus identities and may therefore engage higher-order associational cortical areas, in particular the PFC.

We have previously studied these two levels of sequence processing using a hierarchical “local-global” sequence paradigm where auditory stimuli are presented in short sequences, while measuring macroscopic brain signals. During an entire block of trials, all sequences follow the same pattern, either 4 repeats of the same stimulus (e.g. AAAA), or 3 repeats followed by a deviant (e.g. AAAB). Following the repeated presentation of one of these global sequence patterns, the ability to detect global violations is probed by presenting sequences that deviate from the pattern (AAAB or AAAA, respectively; global deviants). In contrast to the MMN, mismatch responses to such global deviants are delayed, require consciousness, and predominantly arise from higher-order cortical areas of humans and macaques (Bekinschtein et al., 2009; Chao et al., 2018; El Karoui et al., 2015; Uhrig et al., 2014), including the ventrolateral prefrontal cortex (vlPFC). Intracranial recordings of global field potentials indicate that PFC sends feedback signals to upstream cortical areas after the violation of global sequence expectations (Chao et al., 2018). From the observation of such mismatch signals, it was indirectly inferred that the PFC and associated high-level areas comprise an internal mental model of the ongoing sequences. We here demonstrated that neural populations in the vlPFC indeed encoded abstract, global sequence violations.

A central hypothesis of Bayesian predictive processing models is a functional segregation of prediction and error representations, namely different neuronal populations representing internal models or expectations about sensory input and their update, respectively (Bastos et al., 2012; Friston, 2010, 2018; Rao and Ballard, 1999; Walsh et al., 2020). Our results confirm the existence of both error (mismatch) and expectation (or prediction) signals in PFC, as predicted by the theory and in agreement with macroscopic signals recorded with EcoG in the primate brain (Chao et al., 2018; Dürschmid et al., 2019). However, we traced global sequence context and its mismatches to the same population subspace, indicating an integrative level of predictive processing in the PFC.

Frontal cortical areas have also been shown to provide contextual information in the rodent V1 (Hamm et al., 2021), possibly through gain modulation of neuronal activity (Zhang et al., 2014). In our recordings, contextual and mismatch signals were coded by activity vectors intermingled within the same neural population. Specifically, the prefrontal neural population that emitted abstract responses to pure global deviations from a learned sequence also represented the global context. This convergence of predictive-coding computations at the population level may be a particular feature of PFC neurons, which are known for their role in integrative processes and mixed selectivity properties (Parthasarathy et al., 2017; Rigotti et al., 2013). Interestingly, axons from the anterior cingulate cortex, a medial frontal cortical area, projecting to V1 in the rodent brain were not found to be modulated by deviant stimuli (Hamm et al., 2021). This discrepancy might point to a different role of the primate PFC, where we found responses to deviant stimuli, compared to rodent frontal cortex. Alternatively, this could indicate a functional specialization of primate frontal cortical areas, with the vlPFC more likely to reflect both types of responses (deviant and context). It is worth noting that, in an auditory oddball paradigm, context-dependent mismatch responses were indeed observed in rat PFC (Casado-Román et al., 2020).

### An abstract neural code for stimulus transitions

Here, with our control experiment, we found that the neural subspace that conveyed local deviance (mismatch) responses was also activated for any stimulus transition and that this was independent of image identity. There is an ongoing controversy about the mechanisms of mismatch detection. Such a response could stem from two distinct mechanisms: a higher-order process of prediction error, with an active neural response representing the violation of a top-down prediction (Casado-Román et al., 2020; Chao et al., 2018; Hamm et al., 2021; Parras et al., 2017; Summerfield et al., 2008b), or alternatively, a passive feedforward process of stimulus-specific adaptation (Fishman and Steinschneider, 2012; Kaliukhovich and Vogels, 2014; May and Tiitinen, 2010). Based on our results, we were able to falsify the hypothesis that SSA could explain the observed mismatch detection response, by showing how a rare local deviant (last stimulus in XXXY trials) elicited a stronger population response than an equally rare stimulus that was not preceded by a predictable context (WXYZ trials; Fig. 6). However, each stimulus in an WXYZ sequence still elicited a response of the “deviance population”. As this code generalized across images (Fig. 6B light and dashed gray), it can be interpreted as an abstract code for any stimulus change, thus encoding whether the present image is or is not repeated. The response of this “change detector” population decreased with every transition (Fig. 6A) and, as noted, was weaker for a transition following repetitions (XXXY) than for the fourth transition of a sequence (WXYZ). Hence, the adaptation of neurons that encode abstract change information could serve as a building block to encode other structural variables, such as deviance or number.

### Implications for perceptual inference and conscious perception

It has been suggested that PFC participates in a global neuronal workspace (GNW) critical for the representation of any conscious experience (Dehaene and Changeux, 2011; Dehaene et al., 1998). Indeed, several previous experiments indicate that the contents of visual consciousness can be decoded from prefrontal neuronal activity, and that PFC activity signals whether a stimulus was or was not consciously perceived (Kapoor et al., 2018; Levinson et al., 2021; Panagiotaropoulos et al., 2012; van Vugt et al., 2018). The present results are congruent with the GNW hypothesis since they indicate that, in the awake monkey, PFC contains superimposed neural codes for all of the concrete and abstract features of the perceived visual sequences. Some of these features encode expectations of the upcoming sequence, a finding which fits with prior evidence that PFC anticipatory signals may be critical for detecting perceptual ambiguity and biasing conscious perception through ongoing fluctuations (Dwarakanath et al., 2022; Moutard et al., 2015; van Vugt et al., 2018) or the provision of perceptual hypotheses (Summerfield et al., 2008b; Weilnhammer et al., 2021). Our results showing decoding of context from prefrontal populations indeed suggest a strong influence of expectation in the activity of prefrontal ensembles during conscious visual perception.

## Conclusion

Our results reveal that PFC population activity, even in the absence of overt behavior, spontaneously encodes visual sequences at multiple levels, including abstract codes for serial position and/or task phase and sequence pattern, regardless of the particular images used. In this respect, the present results confirm that a representation of abstract sequence structures such as xxxY (“4 items, the last of which is different”) is accessible to non-human primates (Dehaene et al., 2015). Most importantly, it provides insight into the neural code for sequences, which in the PFC relies on a superimposition of multiple vector subspaces, each representing a specific dimension of the perceived sequence. Finally, we showed how these representations can be updated by low- and high-level deviance events. It remains to be shown whether and how these PFC signals influence downstream or upstream cortical areas. Future studies, recording simultaneously from multiple cortical areas and using laminar recordings, will be needed to probe how inter-areal exchanges contribute to predictive processing.

## Acknowledgements

We thank Julien Lemaitre for providing veterinary care, Abhilash Dwarakanath for his help with the data preprocessing pipeline and Pierre-Louis Bellet for providing the brain graphic of Figure 1. This project/research has received funding from the European Union’s Horizon 2020 Framework Programme for Research and Innovation under the Specific Grant Agreement No. 945539 (Human Brain Project SGA3). TIP is supported from the Templeton World Charity foundation (TWCF0562).

## Author contributions

Conceptualization: M.E.B., B.J., S.D, T.v.K., T.I.P., Experimental design: T.v.K. and T.I.P., Analyses: M.E.B, Data acquisition: M.G., T.I.P, Control experiment design: M.E.B and J.B., Control experiment data collection: J.B., Utah array implantation: M.G., B.J. and T.I.P., Writing: M.E.B, T.v.K, S.D. and T.I.P. Visualization: M.E.B

## METHODS

### Lead contact

Further information and requests for resources should be directed to and will be fulfilled by the lead contact, Dr. Theofanis I. Panagiotaropoulos.

### Data and code availability

Code used to generate results is available at https://github.com/mebellet/PrefrontalSequenceModel and data will be made publicly available upon publication.

### Experimental model and subject details

Two adult rhesus macaques (A and H, 9 - 10 kg, 19 and 16 years old, respectively) participated in this study. Both animals were pair-housed. They had previously been implanted with a custom made skull-form-specific titanium headpost and trained on a passive visual fixation task with liquid reward in a primate chair. Daily water access was controlled during the experimental period. All procedures were conducted in accordance with the European convention for animal care (86-406) and the National Institutes of Health’s Guide for the Care and Use of Laboratory Animals. Animal studies were approved by the institutional Ethical Committee (CETEA protocol #A18_028).

### Sequence paradigm

We used an adaptation of the local-global paradigm (Bekinschtein et al., 2009) with visual stimuli. The stimuli were 10 colored images of objects, matched in luminance (Fig. 1D). Fixed pairs of images were used in every experiment, here denoted as stimulus A and B. The paradigm consisted in the presentation of binary visual sequences composed of 4 items. Each item was displayed for 300 ms, with an inter-stimulus interval (isi) of 300 ms (stimulus onset asynchrony, SOA - 600ms). The sequence could be one of four: aaaa or bbbb, denoted as xxxx; and aaaB or bbbA, denoted as xxxY, where the capital letter indicates a *local deviant* item. One sequence was presented per trial which were organized in blocks of 200 trials. During each block, one sequence was used as the frequent, global standard sequence which was established during 50 habituation trials at the beginning of the block. 80% of the remaining 150 trials were global standards and 20% were global deviants, which differed in the last position compared to the standard. Each of the four sequence types was used as the global standard sequence in one block. We will denote a global context according to its global standard sequence: xxxx block for aaaa-frequent and bbbb-frequent or xxxY block for aaaB-frequent and bbbA-frequent. We will furthermore denote trials according to their context: xY|xx indicates a trial with a local deviant that occurs in a block of frequent xxxx sequences. The four trial types are thus xx|xx, xY|xY (global standards) and xx|xY, xY|xx (global deviants).

This two-by-two design enabled us to study effects of lower-order (local) and higher-order (global) sequence regularity. Consider for instance a single xxxY sequence: it ends with a local deviant, an image that violates the repeated structure of the previous three images. However, assuming that monkeys quickly detected the global sequence regularity in a block, the same local deviant, occurring within a block of similar xY global standard trials, is predictable and may no longer generate a global surprise (xY|xY, “predicted local deviant”). Conversely, a rare xx trial, which does not violate the local context, may elicit a global surprise when presented among many xY sequences (xx|xY, hereafter called “pure global deviant”).

Each trial started with the display of a black fixation spot (diameter of 0.3 degree) at the center of the screen. After 300 ms of fixation by the monkey, the fixation point disappeared and the sequence was displayed centrally with stimuli at a size of 8 degrees per visual angle. The animals had to maintain the gaze within a window of 8 degrees of visual angle centered on the stimulus. A liquid reward was given for trial completion 100 ms after offset of the last item.

For monkey H, 5 stimulus pairs were used during a total of 10 experiments and for monkey A, 4 stimulus pairs were used during a total of 7 experiments.

### Repetition versus change control experiment

We performed two sessions of an additional experiment with monkey A during which we showed four different types of sequence chunks containing each 4 images. The types of sequences were XXXX, XXXY, XYYZ and WXYZ, where letters indicate any of 948 grayscale images of objects from the Brainscore database (Majaj et al., 2015), randomly changing with each sequence presentation. Sequence types were uniformly distributed across a recording session so that there was no global context. All stimuli could occur in any position and were presented 1 to 5 times. This experiment served to control for stimulus-specific adaptation that could underlie the deviance response to the last stimulus in XXXY trials. As we used a broad range of stimuli, the images used as WXY are expected to be on average as distant from Z than X is from Y in XXXY trials, thereby leading to the same amount of stimulus-specific adaptation for the population responding to Y (in XXXY) as to Z in (WXYZ).

The timing of the stimuli were the same as in the local-global experiment and the reward was given at 100 ms after offset of the last stimulus.

### Implantation of microelectrode arrays

The macaques were tranquilized in their cage by intramuscular injection of ketamine 1000 (3 mg/kg) and dexmedetomidine (0.015 mg/kg). Once in the operating room, they were placed in a stereotactic frame and deeply anaesthetized (assisted respiration) by inhalation of oxygen (20%) and sevoflurane (1.5 -2%). An intravenous catheter was installed for the administration of physiological fluids (NaCl with 5% glucose, 10 ml/kg/h). A steroidal anti-inflammatory drug, the methylprednisolone (solumedrol 1 mg/kg i.m.) or dexamethasone (dexazone 0,5 mg/kg i.m.) is administered to prevent swelling of the cortex. As well as, an antibiotic (cefazoline 50 mg/kg i.m.) and a morphine derivative (buprecare 0.02 mg/kg i.m.). All surgical procedures were performed aseptically, and recordings of heart rate, respiration patterns, blood pressure and body temperature were monitored throughout the surgery.

Macaques were given a methylprednisolone (monkey A, 1 mg/kg) or dexamethasone dose (monkey H, 0.1 mg/kg) the day before the implantation to avoid brain edema. Monkey H received another dose of dexamethasone (0.5 mg/kg) the day of implantation. The implantation of the gas sterilized multielectrode array began with a longitudinal incision in the skin. The skin and underlying muscle was retracted and a craniotomy was performed over the lateral prefrontal cortex using a surgical drill. The bone flap was removed and then a U-shaped opening in the dura mater was made to expose the cortex. Hyperventilation after dura opening was used to reduce intracranial pressure and avoid swelling of the cortex. The Utah microelectrode array was implanted into the inferior convexity of the prefrontal cortex, 1–2 mm anterior to the bank of the arcuate sulcus and below the ventral bank of the principal sulcus, using a pneumatic inserter (Blackrock Microsystems). The dimensions of the array was 4 × 4 mm in a 10 × 10 electrode configuration resulting in an electrode-to-electrode distance of 400 μm. The electrode length was 1 mm. The titanium connector that can be connected to the electrophysiological recording device was implanted on the skull with titanium screws. Then the dura mater was sewn back together, the bone flap was reinserted and secured by a thin titanium strip. Finally, the skin was sutured. After the electrode array implantation, injections of antibiotics (cefazoline 50 mg/kg i.m.) were given for 10 days and buprenorphine (0,015 mg/kg i.m.) for 3-5 days depending on the pain level.

### Data preprocessing

The recorded broadband signals were preprocessed using MATLAB (MathWorks). Broadband neural signals (0.1 - 30 kHz) were recorded with a Cerebus neural signal processor system (Blackrock Microsystems) and bandpass filtered offline between 0.6 - 3 kHz using a 2nd order Butterworth filter. Spikes were detected with an amplitude threshold set at five times the median absolute deviation and spike events larger than 50 times the mean absolute deviation were discarded. Further, spike events with an inter-spike interval of less than the refractory period of 0.5 ms were also discarded. Spike times were aligned to the onset of the photodiode signal indicating the actual time of presentation of the last item in a sequence.

All further analyses were performed with Python. Firing rates of individual sites were computed from the spike times in non-overlapping bins of 25 ms and smoothed with a gaussian kernel corresponding to 50 ms standard deviation. For the data shown in Figure 6 and Suppl. Figure 5, firing rates were computed with a moving average window of 50 ms and a step size of 10 ms in order to obtain a better temporal resolution.

### Single channel analyses

To quantify the modulation of single channel spiking activity by local and global deviance, we only considered recording sites that were significantly modulated by the task. As a criterion for task modulation, we tested if there was a difference in firing rate during the 300 ms fixation period prior to sequence onset and the first 300 ms after presentation of the last sequence stimulus. We used pairwise two-tailed *t*-test per recording site and recording session and false discovery rate (FDR; Benjamini and Hochberg, 1995) across all sites and sessions within an animal to correct for multiple comparisons. Sites with a corrected p-value <= 0.05 were regarded as being modulated by the sequence task. We tested each of the task-modulated sites for effects of local and global deviance, using a Mann-Whitney-U test (Wilcoxon, 1992) on the average firing rate during 1 sec following the onset of the last stimulus, thus taking into account non-normal distribution of firing rates and substantially differing sample sizes in case of global deviants. For the summary of the results across recording sessions, we used FDR across tested sites and sessions per animal. A site with a corrected p-value <= 0.05 was regarded as being modulated by the respective variable. For the scatter plot in Fig. 2A, FDR was not applied and colors indicate sites that have an uncorrected p-value of below 0.05, either for the global effect (cyan) or local effect (orange) only or independently for both effects (dark gray).

### Population analyses

#### Multivariate linear regression

We assessed how the sequences were represented on the neural population level by computing the axes across the MUA space that carried most information about the variables *stimulus identity*, *global context*, *local* and *global deviance*, without pre-selection of recording sites. For this, we used multivariate linear regression, as in the subspace analysis in (Mante et al., 2013). We performed two separate analyses, one to study the representation of the sequences prior to the last stimulus, and one for the time after onset of the last stimulus, in order to measure responses to deviants. For the time before the last stimulus onset, the variables stimulus identity and global context were considered, whereas the trial condition after onset of the last stimulus was defined by the variables stimulus identity, local and global deviance. As deviance responses following the last stimulus might be dynamic, we performed a separate regression per time bin between 0 until 1.4 sec after onset of the last stimulus.

The multivariate linear regression was performed separately for each recording channel with the above-mentioned sequence variables as independent variables and the MUA (*r*) of channel *i* (in a time bin *t*) as dependent variable:

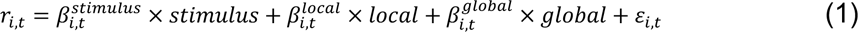

The above equation holds for time bins *t* after presentation of the last stimulus. For the analysis of sequence structure representation before the last stimulus, the responses between 0 - 1.8 sec after sequence onset were averaged per trial and a single regression was performed per recording channel.

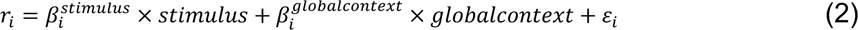

ε is a noise parameter per channel (and time bin). *r* is a vector of dimension *Ntrials*, as are the independent variables stimulus, local, global and global context. Those were dummy variables, with A = -1, B = 1 for the stimulus variable; local or global standards = -1, local or global deviants = 1; xx block = -1 and xY block = 1. This approach results in a coefficient *β* per channel, variable, (and time bin) that indicates how much the firing of a channel was influenced by a certain variable.

The set of the 96 coefficients across all channels for one sequence variable *k* (and time point *t*) constitutes a 96-dimensional vector ^(k)^ (or ^(k)^t) that we denote as the population axis representing this sequence variable. Note that Mante et al. 2013 orthogonalized these axes and denoised them using principal component analysis. We chose not to add these steps after regression in order to measure orthogonality resulting directly from the regression and because the data did not require further denoising.

#### Decoding from the population trajectories

In order to use the resulting population axes for decoding, the MUA of all channels was projected onto the population axis of each sequence variable *k*, respectively.

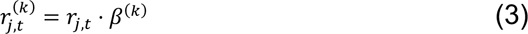

or

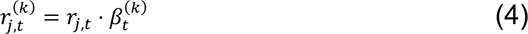

for time-varying population axes.

*j* is the trial index. rj is a 96-dimensional vector of the population firing rate in a single trial *j* and time bin *t*. rj,t is a scalar and corresponds to the dimensionality-reduced population activity in one trial *j* and time bin *t* in the subspace carrying most information about a sequence variable *k*. This trial-by-trial projection was then used to classify trials according to each sequence variable. The sign of these projections was dependent on the definition of the independent variables (see above). A positive activation along the axis coding of stimulus identity, e.g., corresponded to stimulus B, whereas a negative activation corresponded to stimulus A. As a measure of decoding performance, we computed the area under the ROC curve (AUROC) by varying the decision boundary for classification.

#### Cross-validation

The decoding performance was cross-validated both within and across sessions. Within each session, we used 10-fold cross-validation, meaning that 90% of all data was included for the regression and the remaining 10% for projecting test data onto the obtained axes. This was repeated 10 times so that all trials were used for testing. We shuffled the data prior splitting and ensured a balance of trial conditions in the training data. The reported performance within a session is the AUROC across all tested trials.

For the cross-validation across sessions for the variables *global context, local and global deviance*, we used the population axes from one random training fold of each session and projected all trials from all other sessions onto these axes. We then computed the AUROCs for each pair of training and test sessions and reported the performance separately for pairs that had the same or different stimulus pairs.

### Decoding of serial position

We used multinomial logistic regression to predict the item position in a sequence based on the neural population responses. This was a classification with 4 target classes (item 1-4). The 300 ms after onset of a stimulus, shifted by 100 ms, were labelled with the item number of the most recent stimulus. The activity of each channel was averaged in these intervals, resulting in 4 values per trial. The 4 items from all trials were pooled and used to train the classifiers, using 10-fold cross-validation. Only xx trials were used for training. For testing, the activity in each test trial and time bin between 100 ms prior to sequence onset until 1.4 sec after sequence offset was passed through the trained classifier. This resulted in predictive probabilities for item 1 to 4 over time and allowed us to study the dynamic encoding of item position throughout a trial. We also assessed item position classification on incomplete trials. The monkeys could break fixation at any time during a trial by moving the gaze outside of the 6 degree fixation window which aborted the presentation of the sequence. We computed the predictions for item position 1 to 4 for trials interrupted after presentation of the first, second or third stimulus. Note that the fixation break could have occurred at any time between onset of one stimulus and onset of the next stimulus, meaning that the time the monkeys perceived the last stimulus varied within one condition.

### Cross-condition decoding of global deviance

To test for encoding of global deviance irrespective of local deviance, we trained a separate binary classifier to predict global deviance for the time after last stimulus onset, on xx trials only and tested on xY trials. We used logistic regression on the pooled data from all sessions, per animal. This was done to reduce the impact of the block structure of the task within each session, which could have been problematic in this case, as the classifier was trained from global deviants and standards from separate blocks (e.g. rare xx in an xY block vs. frequent xx in an xx block). Decoding performance was again measured as AUROC for each time bin, separately for each session.

### Decoding of deviance or change in control data

We used logistic regression to predict whether a sequence chunk was XXXY or XXXX (deviance decoder), based on the activity after the last stimulus. We used a time-varying decoder in time bins of 50 ms, and with a step size of 10 ms. Image identities were balances in both conditions, i.e. we only included stimuli for training that occured in XXXY and XXXX chunks, resulting in 757 unique images. Decoder performance was cross-validated by leaving trials with one image out for testing. We hence trained 757 different classifiers. Images that did not occur in both conditions (191 different images) were only used once for testing but not included in the training set.

We additionally trained a decoder to detect any change from a repetition by contrasting the response to alternations and repeats in the second and third position of sequence chunks (i.e. the second stimulus in WXYZ and XYYZ trials vs. the second stimulus in XXXY trials as well as the third stimulus in WXYZ vs. the third stimulus in XYYZ trials). The same cross-validation approach was used to test this decoder.

### Assessment of learning effects during the habituation period

We measured whether the code for global context evolved over the course of the habituation period. For this, we projected the activity of single habituation trials (0 - 1.6 sec after sequence onset) onto the population vector that separated xx from xY blocks during the test trials within the same session. As we had used 10-fold cross-validation to obtain those vectors, we also obtained 10 projections of the same habituation trials. We averaged those projections across folds to obtain one activation value for the global context trajectory per trial. To assess learning, we tested the difference in the activation during the first trial vs. the 50th trial using a paired t-test with N=14 blocks in monkey A and N=20 blocks in monkey H. This was done separately for xx and xY blocks, assuming that xx blocks would show an evolution towards a more negative activation (which was defined as the xx block direction) and xY blocks an evolution towards a more position activation (xY block direction).

### Statistics

Statistical tests for single-channel effects are described in the paragraph “Single channel analyses”.

#### Random permutation test

To test the significance of decoding performance from population trajectories, we used a random permutation test with cluster-based correction for multiple comparisons (Maris and Oostenveld, 2007). After estimating the population axes and projecting single trials onto these axes, we generated 100 surrogate datasets by shuffling the trial conditions of test trials. We then computed the AUROCs for the different sequence variables based on the trajectories with the shuffled trial labels. We averaged the true AUROCs across recording sessions (10 in monkey H, 7 in monkey A) and likewise obtained 100 surrogate session-averages. The true AUROCs per sequence variable were transformed into t-values by subtracting the average over the permutations and dividing by their standard deviation, separately for each time point. Absolute t-scores that passed a threshold of 3 standard deviations were candidates for significant clusters in time. This procedure corresponds to a two-tailed test. A correction for multiple comparisons across time was performed by comparing the sum of t-values within each true cluster with the sum of t-values within surrogate clusters. Those surrogate clusters were obtained by transforming each of the 100 permutation samples into t-values by subtracting the mean of the remaining 99 samples and dividing by their standard deviation. If a true cluster had a sum of absolute t-values larger than 95% of the largest surrogate clusters, it passed the threshold for significance which was set to a type-1 error of 5%. For the test of decoding performance across sessions, the same procedure was followed. First, we averaged for each test session the performance based on the decoder trained on the different possible training sessions (same or different stimulus pair). Second, we averaged the 10 or 7 test sessions. The same was done for the surrogate AUROCs based on shuffled trial labels.

The same test was also performed for the cross-condition decoding of global deviance and the effect of deviance onto eye movements (see below).

#### Analysis of eye movements

The eye velocity (v) was measured from the non-calibrated horizontal (x) and vertical (y) eye position recording by computing the difference between time bins (t).

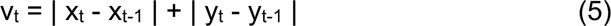

To test for an effect of local or global novelty on eye movements, the median smoothed velocity (20 ms moving average) in each condition was computed across trials from all sessions and tested using a random permutation test (see above). We then controlled for eye movements in the time window during which there was a significant effect of deviants, +-100 ms to be more inclusive, by removing deviant trials with an average velocity in this time period above the median eye velocity in the standard trials. The effect of deviance in the neural data was visualized for all trials vs. the controlled case (Fig. S5).

---

**Figure S1.**
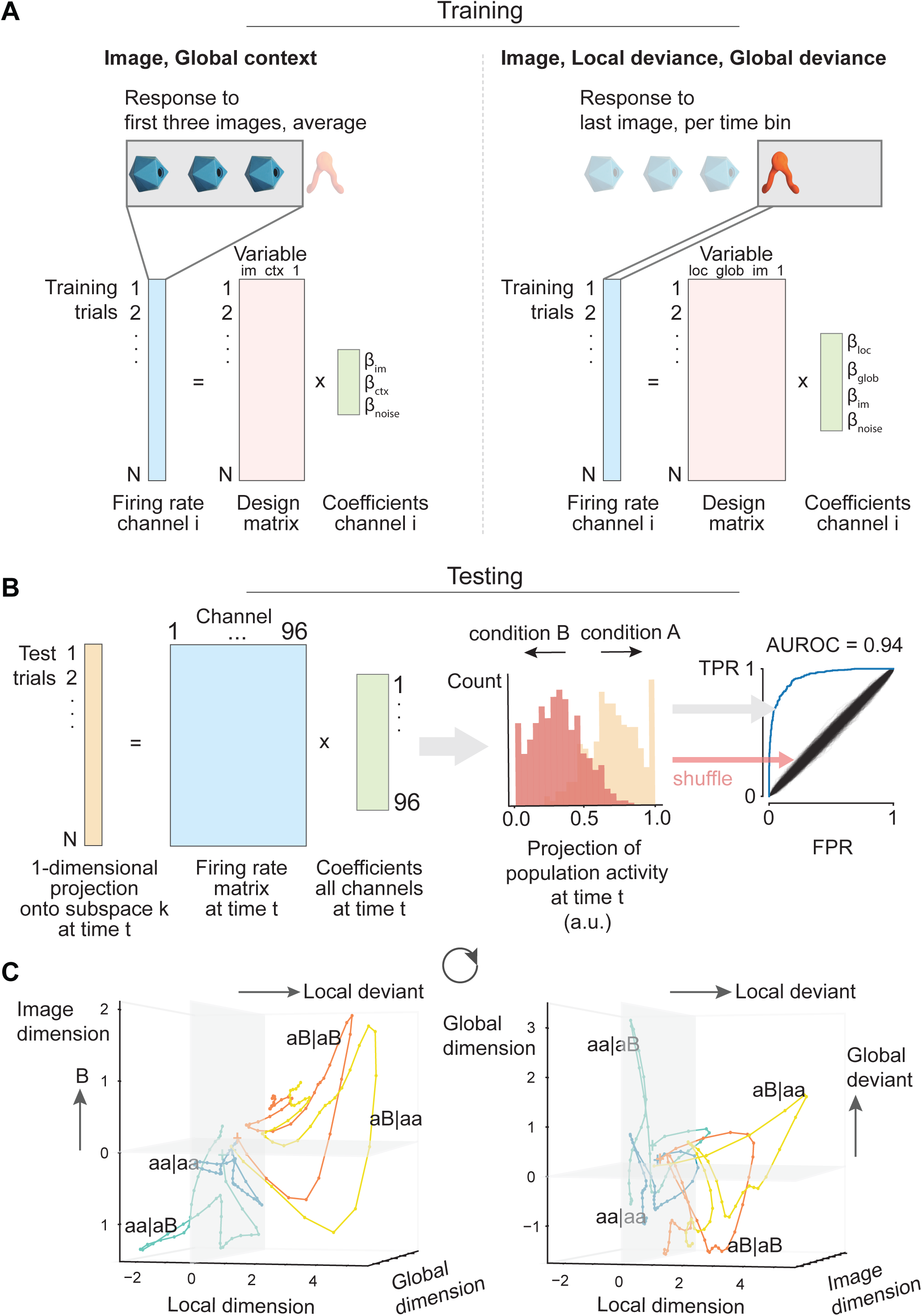
Illustration of the neural population analysis. Related to Figure 3. **A:** Multivariate linear regression is used to determine coefficients per channel and variable. Included variables were image identity (im) and global context (ctx) on the right and image identity, local deviance and global deviance for the time-varying decoder on the left. **B**: For testing the decoder, the neural population activity in each time bin was projected onto the coefficient vector of all channels (left). The projection of test trials (middle) was used to compute the ROC curve for each variable of interest (right) and trials were shuffled to test statistical significance of the resulting AUROC (black lines show shuffle ROCs). FPR = false positive rate, TPR = true positive rate. **C**: Trial-averaged projection onto subspaces for different sequences revealing specific trajectories along each subspace. The time between 0 and 1 sec after onset of the last stimulus is shown.

**Figure S2.**
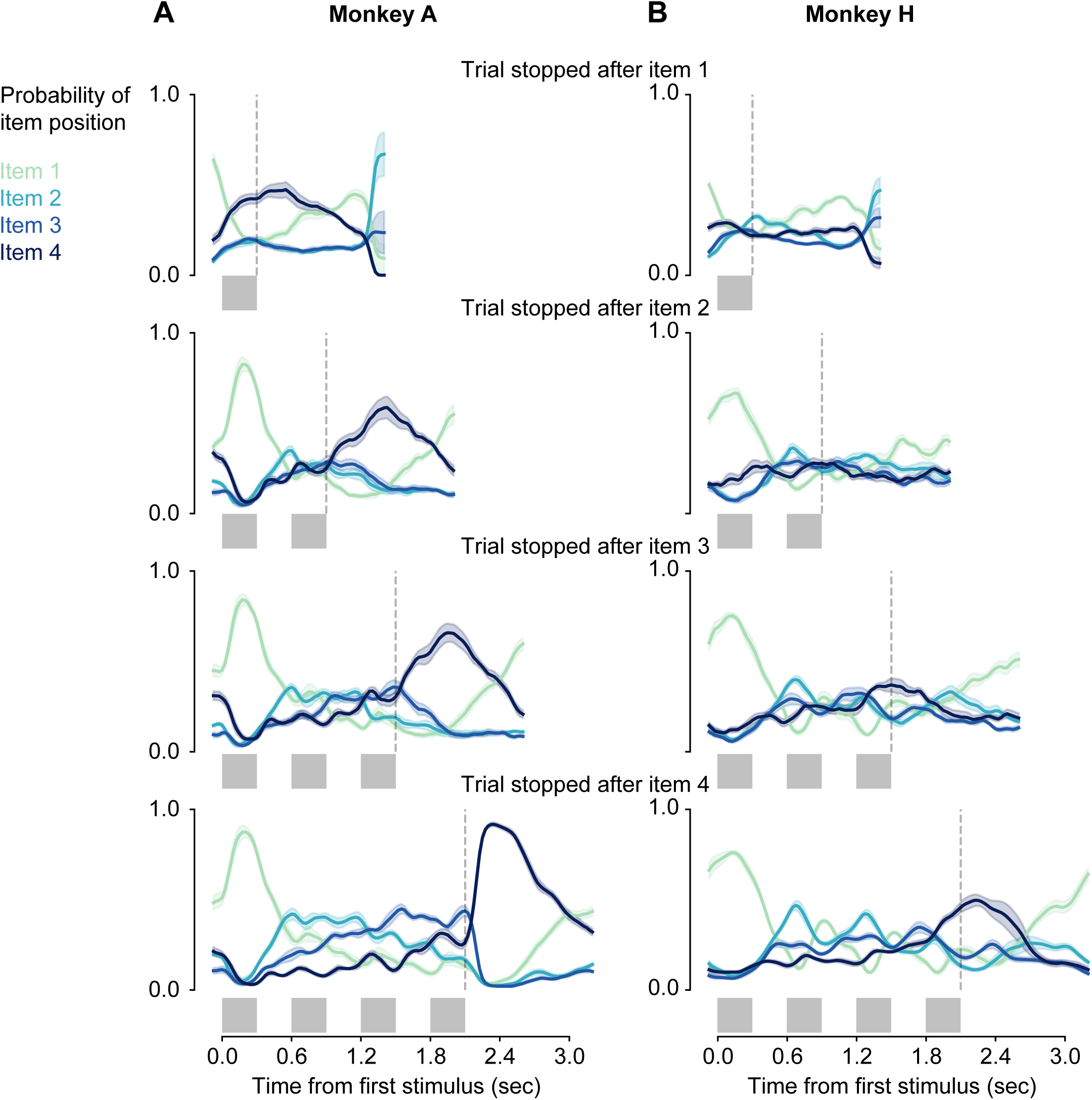
Decoding of sequence position in incomplete trials. Related to Figure 3. A linear decoder was trained to predict the position of a stimulus in a sequence (1 out of 4 possible positions) during completed trials. As the timing was fixed for all sequences, this might correspond to a decoder for the phase of a trial or for time. To test if the components encoding position 1-4 reflected time or a combination of task phase and numerical information, we assessed the population activities for incomplete trials. The population response locked to the fourth stimulus in a sequence (dark blue) shifted in time with the break of fixation, indicating that this component was related to the end of a sequence or trial. The component peaking between position 3 and 4 (blue) on regular trials rose towards the end of the regular sequence and dropped after the fixation break. This population response could encode anticipation of trial end or reward or as well reflect a numerical integration of the stimuli until the trial end. The component coding for the first stimulus in a sequence did not peak for the first item in case the monkey stopped fixating during the presentation of the first item, indicating that this activity reflected a visual response to the stimulus, which might not always be seen in case of such early fixation breaks. The findings show that the neural population responses that allowed the read-out of item position were related to the task phase rather than reflecting continuous time. The gray boxes below each graph show the periods of stimulus presentation. The dashed gray lines indicate the latest time point at which the monkey broke fixation. The earliest time point of fixation break was at onset of the last-presented stimulus.

**Figure S3.**
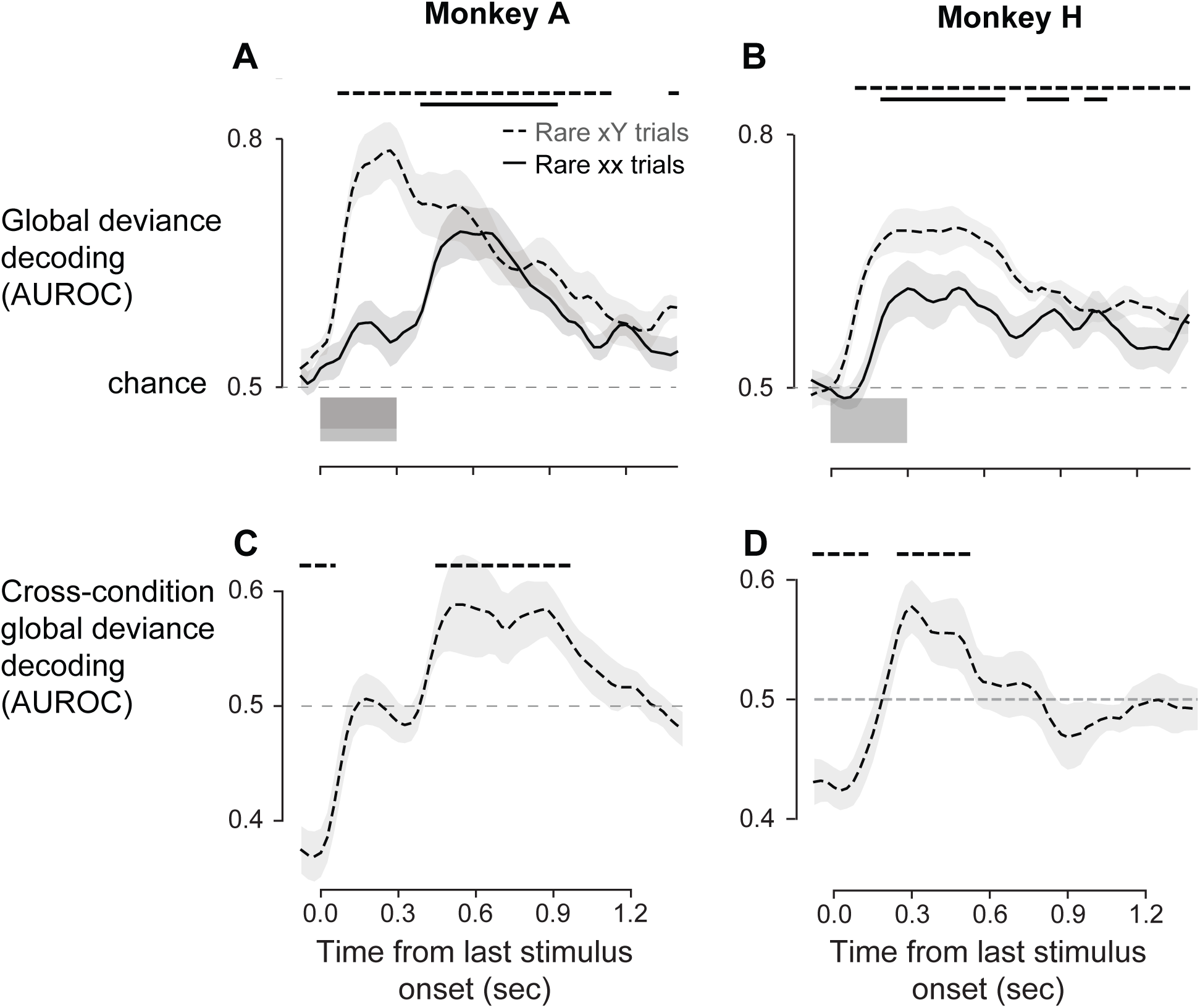
Decoding of global deviance irrespective of local deviance. Related to Figure 3. **A,B**: The time-varying global deviance decoder (from Fig. 3 E and J) was tested separately for rare vs. frequent xY trials (dashed line) and rare vs. frequent xx trials (solid line). Both trial types evoke a global deviance response. The effect in xY trials rises earlier than in xx trials and the deviance response to rare xx trials occurred earlier in monkey H than in monkey A. One reason for this could be that we recorded from functionally slightly distinct neural populations in the two animals. Alternatively, the processing of the global sequence violation in rare xx trials might have required a longer integration time in monkey A. **C,D**: “Cross-condition” global deviance decoding performance. The decoder was trained with rare vs. frequent xx trials only and tested on rare vs. frequent xY trials. A performance above 0.5 indicates a shared global deviance code in both trial types (rare xx and rare xY), whereas a significant performance below 0.5 can be explained by the fact that the comparison between block types is reversed for both trial types (xx|xY vs. xx|xx for training, xY|xx vs. xY|xY for testing). Hence, this corresponds to an effect of global context prior to the global deviance. In all panels, the curves indicate average performance over sessions and shaded areas show +-s.e.m. The horizontal lines on top indicate time periods of significant decoding performance.

**Fgure S4.**
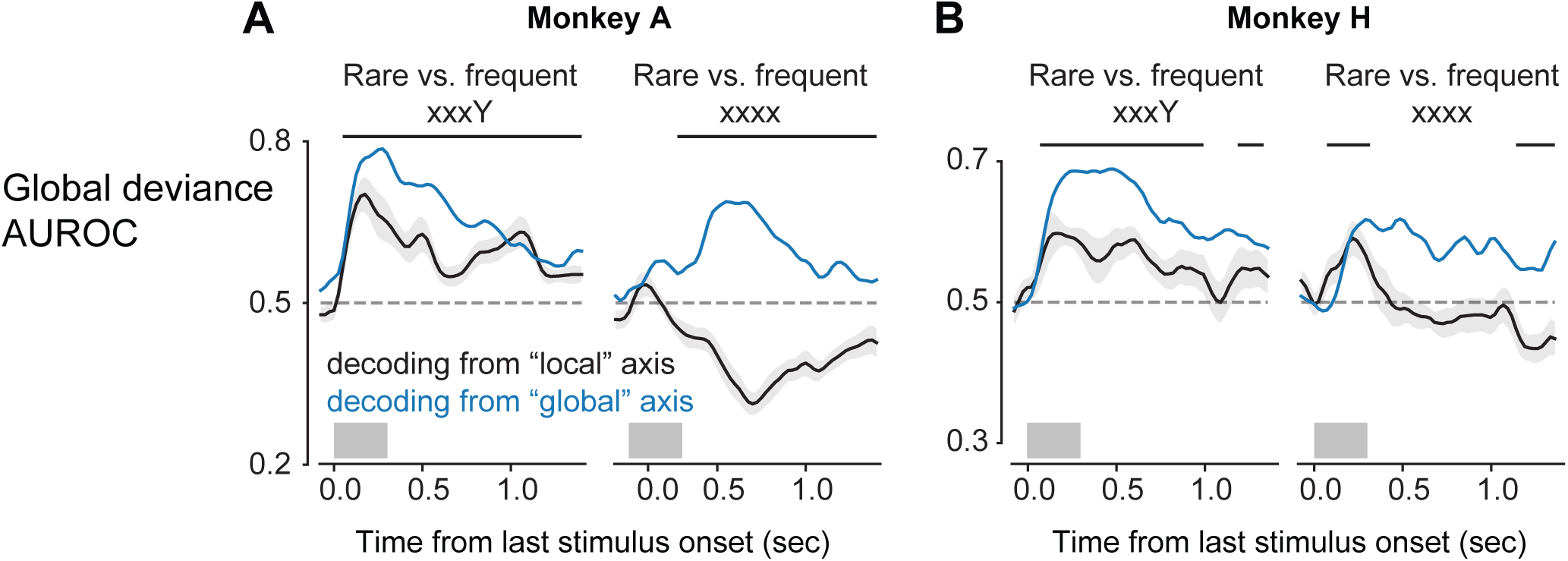
Decoding of global deviance from the neural subpopulation coding for local deviance. Related to Figure 3. The trial-by-trial projections of the MUA onto the population vector encoding local deviants (“local” axis) were used to decode global deviants. This was done separately for rare vs. frequent xxxY and rare vs. frequent xxxx sequences. Black lines show this cross-condition decoding performance averaged over sessions. The blue line serves as a reference and indicates the average decoding performance from the projections onto the population vector coding for global deviance (“global” axis). Decoding performance was measured as area under the ROC curve (AUROC). While rare xY trials could be decoded from the activity along the local axis (**A** and **B left**), rare xx trials led to an opposite effect in this population, indicated by an AUROC below 0.5 (**A** and **B right**), which was significant only in monkey A. This indicates that local and global deviants were encoded by distinct neural subpopulations. Note that in monkey H, rare xx trials could be decoded from the early “local” axis, probably indicating that this early subpopulation carried feedforward adaptation effects present for both global deviant types.

**Figure S5.**
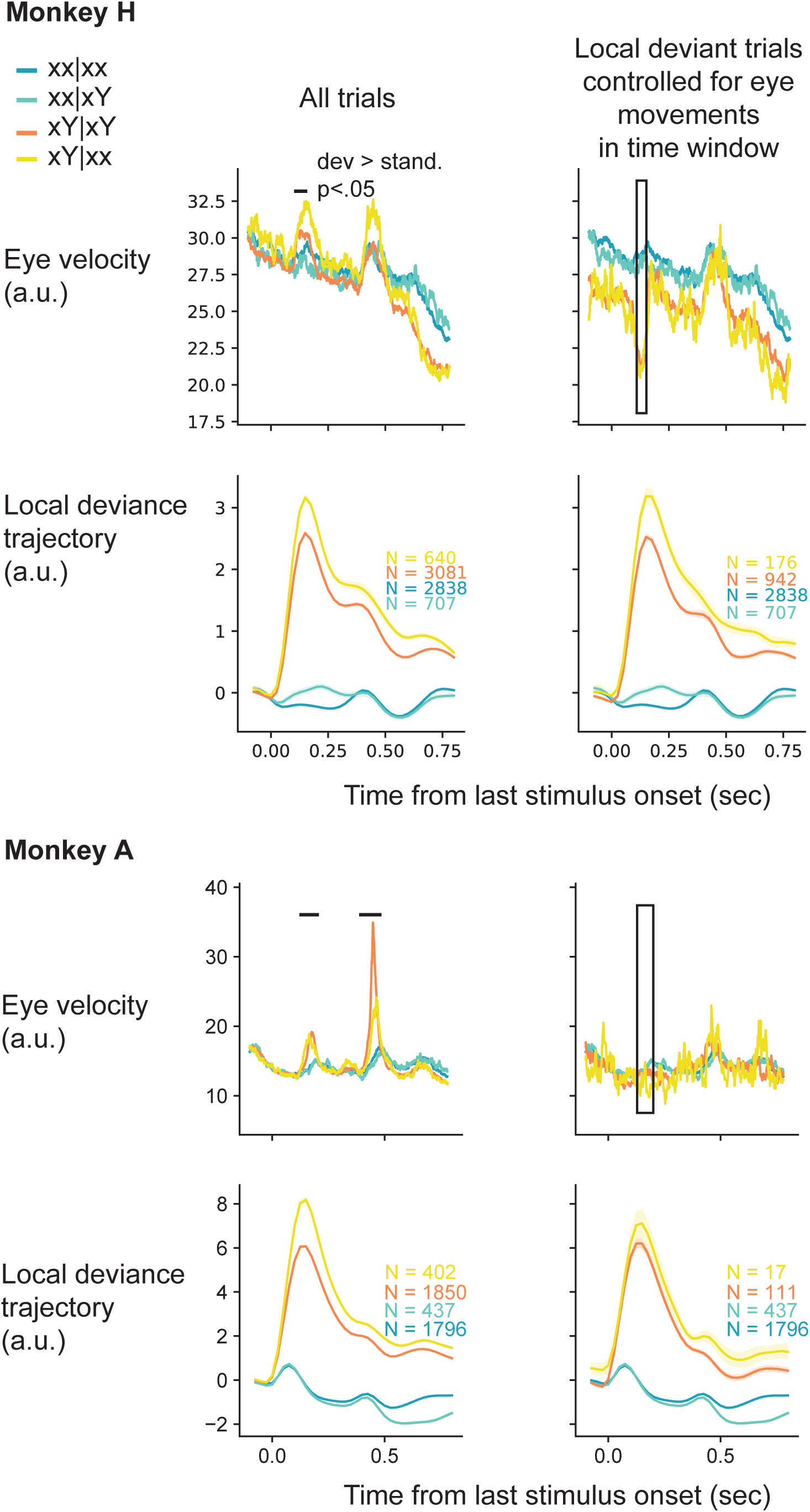
Eye movements do not explain neural deviance signals. Related to Figure 3. The eye velocity was a behavioural read-out of local surprise, indicated by the horizontal black bars on top of the graphs. No main effect of global deviance onto eye movements was found. After removing local deviant trials with an eye velocity above the median eye velocity of local standard trials in the time window indicated by the black box, the effect of local deviance in the neural data remained unchanged (see population trajectories). Graphs show the median eye velocity across all trials and sessions and the average population trajectory of the local deviance subspace across all sessions. Numbers next to the graphs indicate the underlying number of trials.

**Figure S6.**
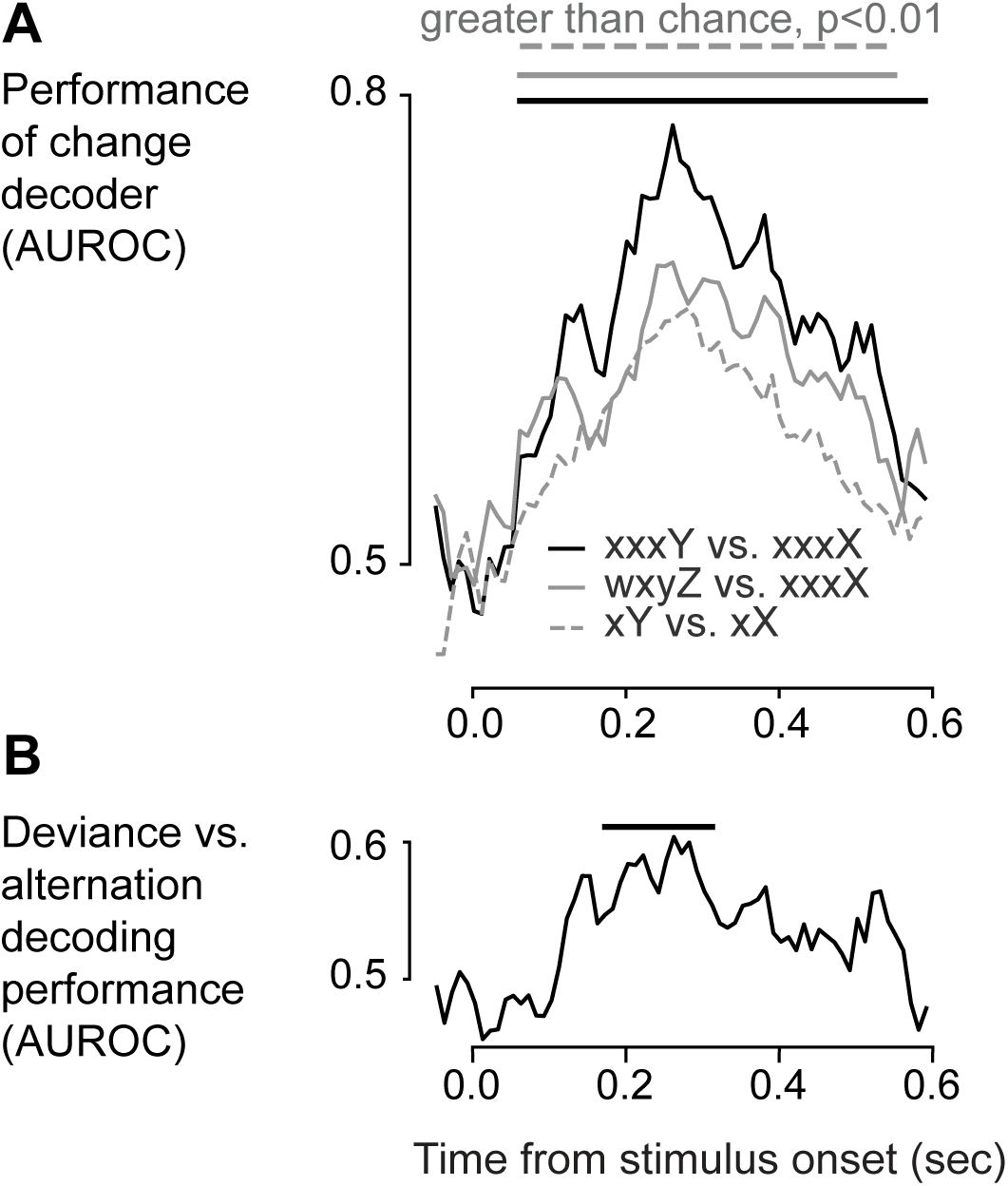
Deviance and change predictions by a decoder trained to detect change. Related to Figure 6. On the data shown in Figure 6, a neural decoder was trained to predict stimulus change, using stimulus transitions vs. repetitions for training (Methods), and validated with leave-one-stimulus-out cross-validation. **A** and **B** show same metrics and tests as Figure 6 B and C but using the output of the change decoder. **B**: Also for this slightly different population, rare stimuli that violate a local pattern of repetitions (XXXY) yielded a significantly higher response compared to any rare stimuli without preceding regularity (WXYZ).

